# Receptor tyrosine kinases activate heterotrimeric G proteins via phosphorylation within the interdomain cleft of Gαi

**DOI:** 10.1101/2020.08.09.229716

**Authors:** Nicholas A. Kalogriopoulos, Inmaculada Lopez-Sanchez, Changsheng Lin, Tony Ngo, Krishna Midde, Suchismita Roy, Nicolas Aznar, Fiona Murray, Mikel Garcia-Marcos, Irina Kufareva, Majid Ghassemian, Pradipta Ghosh

## Abstract

The molecular mechanisms by which receptor tyrosine kinases (RTKs) and heterotrimeric G proteins, two major signaling hubs in eukaryotes, independently relay signals across the plasma membrane have been extensively characterized. How these hubs crosstalk has been a long-standing question, but answers remain elusive. Using linear-ion-trap mass spectrometry in combination with biochemical, cellular, and computational approaches, we unravel a mechanism of activation of heterotrimeric G proteins by RTKs and chart the key steps that mediate such activation. Upon growth factor stimulation, the guanine-nucleotide exchange modulator, GIV, dissociates Gαi•βγ trimers, scaffolds monomeric Gαi with RTKs, and facilitates the phosphorylation on two tyrosines located within the inter-domain cleft of Gαi. Phosphorylation triggers the activation of Gαi and inhibits second messengers (cAMP). Tumor-associated mutants reveal how constitutive activation of this pathway impacts cell’s decision to ‘go’ *vs*. ‘grow’. These insights define a tyrosine-based G protein signaling paradigm and reveal its importance in eukaryotes.

**Significance Statement:** Growth factors and heterotrimeric G proteins are two of the most widely studied signaling pathways in eukaryotes; their crosstalk shapes some of the most fundamental cellular responses in both health and disease. Although mechanisms by which G protein pathways transactivate growth factor RTKs has been well-defined, how the reverse may happen is less understood. This study defines the key steps and cellular consequences of a fundamental mechanism of signal crosstalk that enables RTKs to transactivate heterotrimeric G protein, Gαi. Mutations found in tumors shed light on how derailing this mechanism impacts tumor cell behavior. Thus, findings not only show how cells integrate extracellular signals *via* pathway crosstalk, but also demonstrate the relevance of this pathway in cancers.

## Introduction

The flow of information from external environmental cues to the interior of the cell is controlled by a complex array of proteins at the plasma membrane. Although signal transduction is traditionally studied in a reductionist fashion by dissecting a single pathway/cascade at a time, it is well-known that these distinct signaling pathways crosstalk at multiple levels. Such cross-talks between multiple pathways generate larger complex signaling networks that ultimately control cell fate (1–4).

In eukaryotes, two of the most widely studied signaling pathways are those that are initiated by the receptor tyrosine kinases (RTKs) and by the 7-transmembrane receptors coupled to heterotrimeric G proteins (GPCRs). Upon ligand binding, growth factor RTKs become auto-phosphorylated on their cytoplasmic tails, creating docking sites for the recruitment and phosphorylation of a variety of adaptor proteins that propagate the signal to the cell’s interior (5). Heterotrimeric G proteins, on the other hand, serve as molecular switches, canonically acting downstream of GPCRs (6, 7). Agonist-bound GPCRs act as receptor guanine-nucleotide exchange factors (GEFs) for heterotrimeric G proteins, triggering GDP to GTP exchange on Gα and releasing Gβγ subunits; GTP-bound Gα monomers and Gβγ dimers go on to bind and transduce signals via a variety of effectors (6).

Although it has been suggested that these two pathways crosstalk (8–12), in that G proteins may be activated downstream of RTKs (13–23), whether and how this process takes place in cells and its biological significance remains ambiguous. Published works from the late 1980’s and early 1990’s have suggested that tyrosine phosphorylation of G proteins is one such mechanism (24–27); however, the identity of these sites and how they may impact G protein activity remained unknown. Here we define the key steps of a mechanism utilized by cells to transduce tyrosine-based signals directly from RTKs to trimeric G proteins and demonstrate the cellular consequences of such crosstalk.

## Results

### Growth factor RTKs phosphorylate Gαi

High throughput mass spectrometry studies (HTP-MS) (20, 22, 23, 25, 28, 29) have reported phosphorylation of Gαi on several tyrosine residues (Fig. 1a); some of these cluster around the αF helix and Switch-I loop and are buried within the interdomain cleft (circle; Fig. 1b). To determine whether Gαi undergoes tyrosine phosphorylation, we conducted in-cell kinase assays by immunoprecipitating Gαi3 from serum-starved cells stimulated or not with Epidermal Growth Factor (EGF) (Fig. 1c) or insulin (Fig. 1d) and analyzed them by immunoblotting with an anti-Pan-pTyr antibody. Gαi3 was indeed phosphorylated in response to growth factor stimulation (Fig. 1c-d). To distinguish whether the observed phosphorylation in cells is due to RTKs, or due to non-RTKs such as Src family kinases that are also activated downstream of RTKs, we performed *in vitro* kinase assays using recombinant RTKs and bacterially expressed and purified His-Gαi3 and found that Gαi3 was readily phosphorylated *in vitro* by all RTKs tested, i.e., EGFR, PDGFR and VEGFR (Fig. 1e) and the receptor for NGF, TrkA (***SI Appendix*, Fig. S1a**). Using EGFR as the prototypical RTK, we confirmed that RTKs also phosphorylate Gαi1, Gαi2 and Gαo, but not Gαs (summarized in Fig. 1g; ***SI Appendix*, Fig. S1b**), indicating that α-subunits of the Gi/o subfamily are preferred substrates. By contrast, the non-RTK c-Src efficiently phosphorylated all Gα subunits tested including Gαs (summarized in Fig. 1g; ***SI Appendix*, Fig. S1c**). EGFR, but not Src, showed selectivity for Gαi3 substrate conformation; EGFR preferentially phosphorylated inactive (GDP-bound) Gαi3, while c-Src phosphorylated both inactive and the GTP hydrolysis transition state mimic (GDP+AlF_4_-bound) Gαi3 (Fig. 1f, summarized in Fig. 1h; ***SI Appendix*, Fig. S1d-e**). Noteworthy, c-Src selectively phosphorylated inactive (GDP-bound) Gαs *in vitro* (summarized in Fig. 1h; ***SI Appendix*, Fig. S1e**) consistent with previous work (26, 30, 31). These findings indicate that RTK (EGFR)-dependent phosphorylation of Gαi may be distinct from those that are triggered by non-RTKs (Src): they are similar in terms of selectivity for Gα-subfamilies (Gi/o over Gs) but differ in their preference for nucleotide-dependent conformational constraints (RTKs prefer native and ‘inactive’, over ‘active’ state).

**Fig. 1:**
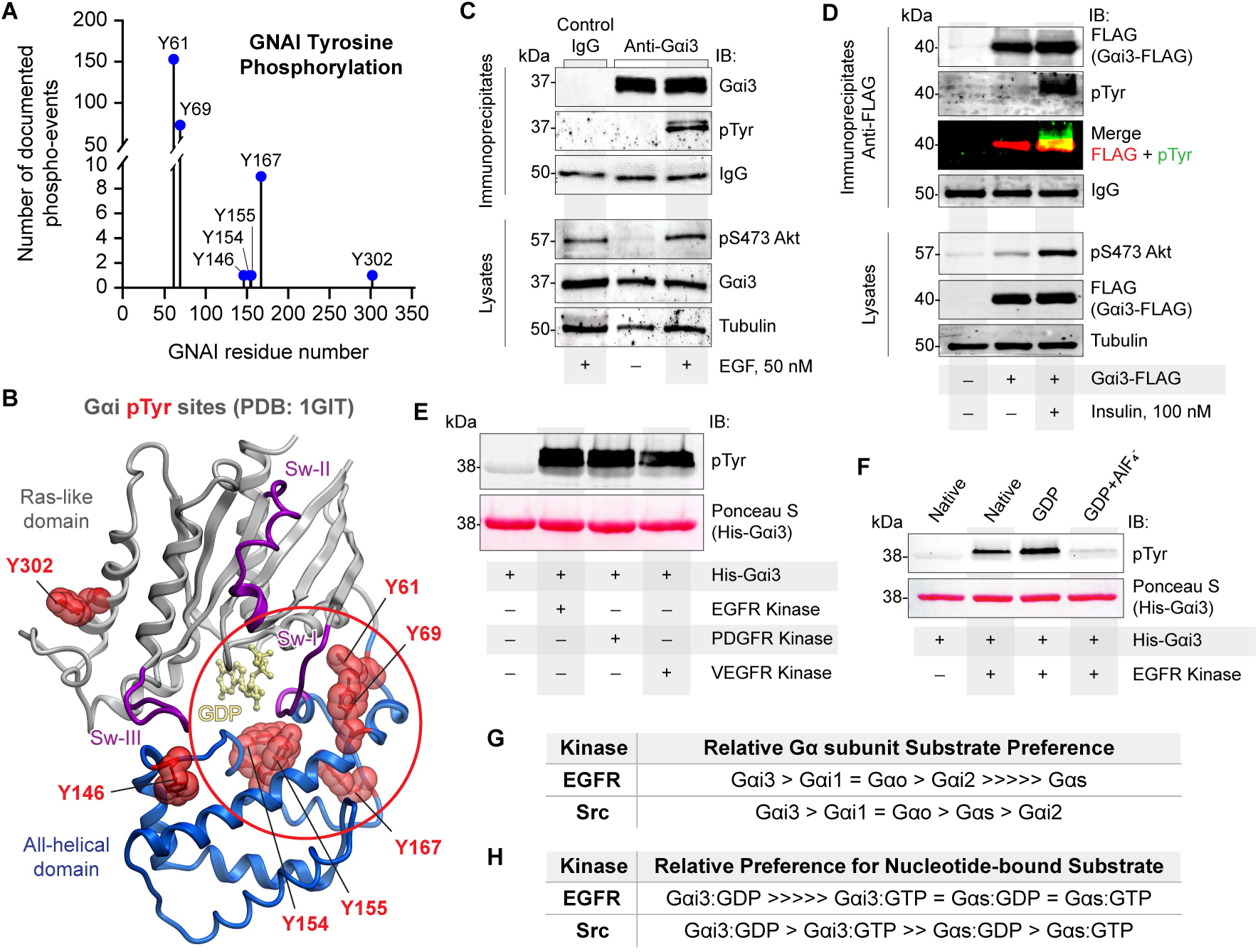
Multiple RTKs directly phosphorylate Gαi. **A**, Lollipop diagram displaying all documented tyrosine phosphorylation events on Gαi1, Gαi2, and Gαi3. **B**, Phosphorylated tyrosines in ‘A’ are projected onto the structure of GDP-bound Gαi1 (PDB 1GIT). Circle highlights several phosphorylated tyrosines that cluster around αF/Sw-I and are bur within the interdomain cleft. **C**, Immunoprecipitation of endogenous Gαi3 from starved or EGF-stimulated HeLa cells. Immunoprecipitates were analyzed for Gαi3 and pTyr by immunoblotting. **D**, Immunoprecipitates of FLAG-tagged Gαi3-WT from starved (-) or insulin-stimulated (+) Cos-7 cells were analyzed for Gαi3 (FLAG) and pTyr by immunoblotting. **E**, *In vitro* kinase assays using His-Gαi3 (2.8 µM) as substrate with multiple recombinant active RTKs (23 nM). **F**, *In vitro* kinase assay using His-Gαi3 (2.8 µM) as substrate in the native, inactive (pre-loaded with GDP), and active state (pre-loaded with GDP + AlF_3_) with recombinant active EGFR (23 nM). **G**, Table summarizing the extent of phosphorylation of various Gαi/o/s subunits observed using recombinant active EGFR and Src kinases [see ***SI Appendix*, Fig. S1b-c**]. **H**, Table summarizing the extent of phosphorylation of various nucleotide-bound Gαi/s subunits observed during *in vitro* kinase assays using recombinant active EGFR and Src kinases [***SI Appendix*, Fig. S1d-e**]. All immunoblots are representative of findings from at least 3 independent repeats.

### Phosphorylation of Gαi in cells requires the cytosolic guanine nucleotide-exchange modulator GIV

Unlike GPCRs, RTKs do not bind G proteins directly, and hence, we asked if phosphorylation of Gαi in cells by RTKs requires scaffolding of the kinase to its substrate. We specifically asked if such phosphorylation requires the large multi-modular protein, GIV (<u>G</u>α-Interacting <u>V</u>esicle associated protein; a.k.a, Girdin), which has been shown by BRET (32) and FRET (33)-based approaches as mediators of the transient formation of ligand-activated RTK•GIV•Gαi ternary complexes. GIV is the prototypical member of a family of cytosolic proteins, guanine-nucleotide exchange modulators (GEMs) (34, 35), which activate Gαi downstream of a myriad of cell surface receptors, including growth factor RTKs, integrins, and GPCRs (36–42). The published structural basis for how GIV scaffolds RTKs to Gαi guided our choice of specific experimental tools (Fig. 2a)-- a Src-homology2 (SH2)-like domain within GIV’s C-terminus recognizes autophosphorylated cytoplasmic tails of EGFR (43) and an upstream 31-aa stretch short motif binds and activates Gαi (44, 45). We chose to use two well-characterized mutations that disrupt the GIV•Gαi interface: Gαi3-W258F (WF) mutant (Fig. 2a-b) which renders the G protein insensitive to the GEF action of GIV (40), and GIV-F1685A (FA) mutant (Fig. 2a, 2c) which can neither bind, nor activate Gαi (41). GIV-SH2-deficient mutants that disrupt the RTK•GIV interface were not considered because they impact receptor autophosphorylation and activation (43). To specifically monitor RTK-dependent phosphorylation, we performed kinase assays in cells in the presence or absence of PP2, a potent and selective inhibitor of the Src-family of non-RTKs (46). In the absence of PP2, EGF triggered the phosphorylation of both Gαi3-WT and the WF mutant, but in the presence of PP2, phosphorylation was only observed in Gαi3-WT (Fig. 2b). These results suggest that the observed phosphorylation in the presence of PP2 (at concentrations that virtually abolished Src activity; Fig. 2b) is likely to be *via* EGFR and demonstrates that EGFR-dependent phosphorylation of Gαi requires GIV•Gαi coupling. Because recombinant EGFR could phosphorylate both Gαi3-WT and W258F (WF) proteins to an equal extent *in vitro* (***SI Appendix*, Fig. S2**), the loss of phosphorylation we observe for the WF mutant in cells is likely due to its inability to come in close proximity to ligand-activated RTKs. Kinase assays in the presence of PP2 on GIV-depleted HeLa cells stably expressing either GIV-WT or GIV-F1685A triggered phosphorylation of Gαi3 only in control and GIV-WT cells, but not in GIV-F1685A cells (Fig. 2c). These findings indicate that GIV•Gαi coupling is a prerequisite for EGF-dependent phosphorylation of Gαi in cells and suggests GIV’s scaffolding action may be one way to create the necessary spatial proximity of the substrate (Gαi) and the kinase (RTK) in cells.

**Fig. 2:**
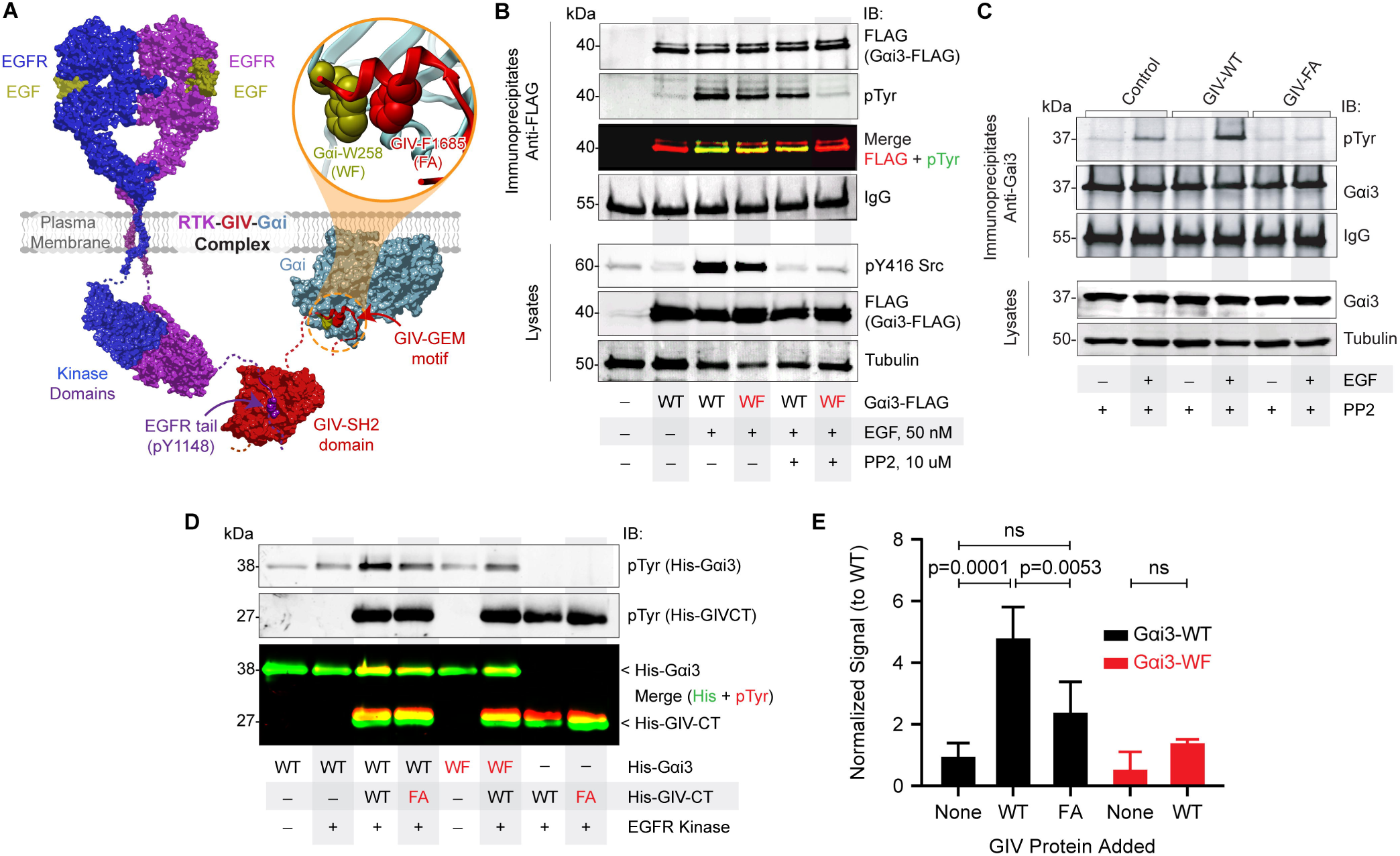
Phosphorylation of Gαi by RTKs requires the cytosolic GEF, GIV. **A**, Schematic of an RTK[EGFR]•GIV•Gαi ternary complex built using a EGFR•GIV homology model (43) and a solved GIV• Gαi structure [PDB 6MHF]. Key residues in the GIV•Gαi interface are highlighted in the inset; mutations used to disrupt the interface are annotated within parentheses. **B**, Immunoprecipitates of FLAG-tagged Gαi3-WT or Gαi3-W258F from starved or EGF-stimulated Cos-7 cells treated or not with the src inhibitor, PP2 were analyzed for Gαi3 (FLAG) and pTyr by immunoblotting. **C**, Immunoprecipitation of endogenous Gαi3 from EGF-stimulated HeLa control, GIV-WT or GIV-F1685A expressing cells. Immunoprecipitates were analyzed for Gαi3 and pTyr by immunoblotting. **D**, Representative *in vitro* kinase assays using recombinant active EGFR (23 nM), His-Gαi3 (937 nM) and His-GIV-CT (1.58 μM) and mutants (Gαi3-W258F, WF and GIV-CT-F1685A, FA; both inhibit GIV’s ability to activate Gαi3) carried out in the presence of GTP to favor nucleotide exchange. **E**, Bar graph displaying quantification of the *in vitro* kinase assays from (**D**). Error bars, ±S.E.M.; n = 4. All immunoblots are representative of findings from at least 3 independent repeats.

Because several phosphotyrosines reported in HTP MS studies were buried in the interdomain cleft (circle; Fig. 1b), we asked if RTK-dependent phosphorylation of Gαi was augmented under conditions that permit conformational plasticity during nucleotide exchange. To this end, we carried out *in vitro* kinase assays in the presence of excess hydrolysable GTP and with the G protein pre-bound to GIV-GEF. The Gαi-W258F and the GIV-F1685A mutants that impair GIV•Gαi coupling were used as negative controls. In the presence of GIV and GTP, EGFR phosphorylated Gαi to greater extent, but only when the GIV•Gαi interaction was intact (Fig. 2d-e). These in-cell and *in vitro* studies using two specific mutants that abrogate GIV’s ability to *bind* and *activate* Gαi show that EGFR-dependent phosphorylation of Gαi requires the scaffolding action of GIV and is facilitated during GIV-dependent nucleotide exchange. The latter is perhaps a consequence of opening of the interdomain cleft to solvent (and thereby, access to the buried tyrosines) during such exchange (45).

### RTKs phosphorylate Gαi on unique sites within the inter-domain cleft

To determine which tyrosine residues are phosphorylated by RTKs, we used linear-ion-trap mass spectrometry (47) and analyzed Gαi3 that was phosphorylated in cells after EGF stimulation or *in vitro* by recombinant EGFR (***SI Appendix*, Fig. S3a**).Three independent analyses were performed in 2 different facilities; none of the samples were subjected to phosphoenrichment. Three phosphotyrosines (pY154, pY155, pY320) were detected (Fig. 3a-d; ***SI Appendix*, Fig. S3a; Dataset S1**); their stoichiometry in cells, as determined by the ratio of phospho- to the total peptides of any given sequence, was ~65% for peptides dually phosphorylated at Y154 and Y155 (pYpY), ~10% for peptides with phosphorylation at Y154 alone; phosphorylation at Y320 was seen in <1% of the peptides; peptides phosphorylated exclusively at Y155 were not detected (Fig. 3d). The same three phosphosites were detected also in insulin-stimulated samples, indicating that these phosphoevents may also be triggered by other growth factors/RTKs besides EGF/EGFR. In the case of the *in vitro* phosphorylated samples of Gαi3 that was pre-loaded with GDP, phosphorylation was detected at the same three sites but to a much lesser compared to that observed in cells (0.25% for dual pY154/pY155, 1.3% for single pY154 and none for pY155 alone; ***SI Appendix*, Dataset S2**), consistent with our prior observation (Fig 2d-e) phosphorylation requires conformational plasticity. Sequence alignment showed that the various Gα-subunits that bind GIV all have Y154, Y155 and Y320 conserved, except for Gαs where the Y at 155 is a Phe(F) (Fig. 3c). We noted two of these sites (Y154/155) were the same buried sites previously detected in HTP studies (Fig. 1a-b); located the αE-helix (aa 151-163), these sites make hydrogen(H)-bonds with the αF-helix (aa 170-177; Fig. 3b-c) within what is dubbed as ‘*the interdomain cleft’* of the Gα-subunit. Using non-phosphorylatable Y→F mutants of Gαi in *in vitro* (Fig. 3e) and in cell (Fig. 3f-g) kinase assays, we confirmed these to be the major sites for RTK phosphorylation because phosphorylation was significantly diminished in the triple tyrosine mutant Gαi3-Y154/155/320F (Gαi3-3YF; Fig. 3e-g), and to various degrees in the individual YF mutants (Fig. 3e). Furthermore, using a custom pYpY-Gαi antibody [raised against a dually phosphorylated peptide derived from pY154/pY155 Gαi3 sequence] and either Phos-tag^®^ SDS-PAGE (Fig. 3h) that resolve phosphoproteins (48, 49) or conventional gels (Fig. 3i), we could detect phosphorylation of Gαi3-WT after an *in vitro* kinase assay with EGFR. These forms were virtually abolished when either Y154 or Y155 was mutated, but unaffected by the Y320F mutation (Fig. 3h). These phosphoforms were weakly detected using the pYpY-Gαi antibody when c-Src-phosphorylated Gαi3 WT or mutant proteins were resolved by Phos-tag^®^ SDS-PAGE (***SI Appendix*, Fig. S3b**), indicating that Src may not phosphorylate Y154 and Y155 as well as EGFR. Defective phosphorylation observed in the case of Y154F, Y155F or the 3YF mutants was not due to catastrophic defects in protein stability, folding and/or functionality because all of them were capable of binding nucleotides and adopting an active conformation as determined by a limited proteolysis assay (***SI Appendix*, Fig. S3c-e**). The ability of c-Src to phosphorylate these Gαi3 mutants as efficiently as Gαi3-WT, as determined by pan-pTyr antibody (***SI Appendix*, Fig. S3f**), confirms our suspicion that RTKs and non-RTKs may both phosphorylate G i3 and that the sites on Gαi3 they preferentially phosphorylate are mostly distinct. Regarding Y154 and Y155, the two tyrosines within *the interdomain cleft* of the GTPase, RTKs preferentially target those sites over Src.

**Fig. 3:**
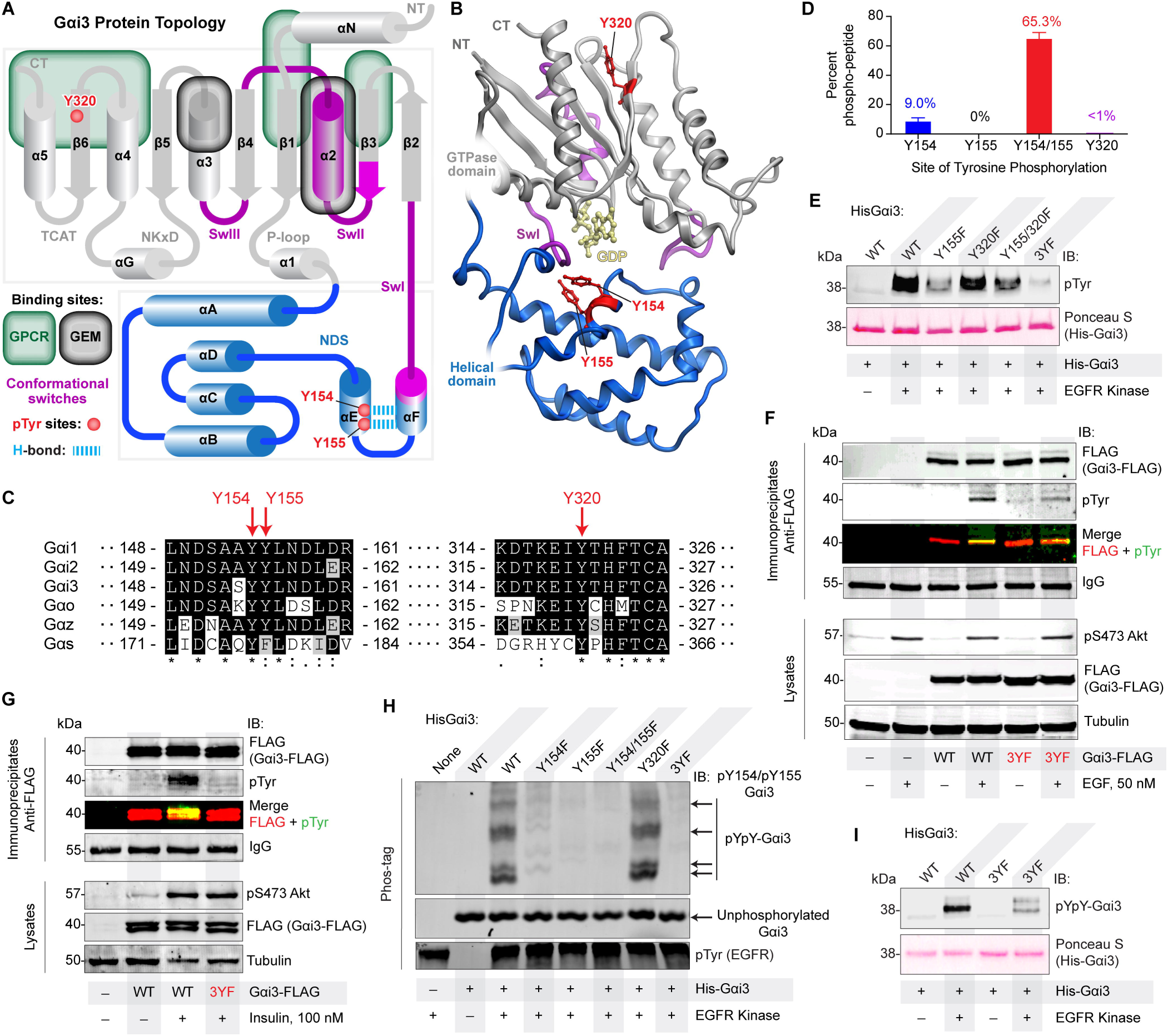
RTKs directly phosphorylate Gαi on Y154, 155, and Y320. **A-B**, The location of tyrosines on Gαi that are targeted by RTKs are projected on a topology map of the Gαi protein (modified from (44)) with conformational switches and binding sites of key interactors marked (A) and the solved crystal structure of Gαi3•GIV co-complex (PDB 6MHF; **B**). **C**, Protein sequence alignment of all Gα-subunits previously shown (86) to bind GIV. Phosphorylation sites identified in this work employing QTRAP5500-assisted phosphoproteomic analyses (see ***SI Appendix*, Fig. S3a**) of *in vitro* and in-cell kinase assays are marked (red arrows). The complete catalog of phosphopeptides can be found in **Datasets S1** and **S2**. **D**, Bar graphs display the ratio of tyrosine-phosphorylated over total peptides [EYQLNDSASY^154^Y^155^LNDLDR and EVY^320^THFTCATDTK] detected in cells (n = 3). Samples were not subjected to phosphoenrichment prior to Mass Spectrometry. Error bars, ±S.E.M.; n = 3. **E**, *In vitro* kinase assays using recombinant active EGFR kinase (23 nM) and either WT His-Gαi3 (2.8 µM) or various non-phosphorylatable YF mutants as substrate. **F-G**, Immunoprecipitates of FLAG-tagged Gαi3-WT or Gαi3-3YF from EGF (**E**)- or Insulin (**F**)-stimulated Cos-7 cells treated with the src inhibitor, PP2 were analyzed for Gαi3 (FLAG) and pTyr by immunoblotting. **H**, *In vitro* kinase assays using recombinant active EGFR kinase (23 nM) and either WT His-Gαi3 (937 nM) or various non-phosphorylatable YF mutants run on Phos-tag^®^ gel and immunoblotted wit custom rabbit polyclonal anti-pYpY-Gαi3 antibody. **I**, *In vitro* kinase assays using recombinant active EGFR kinase (23 nM) and either WT His-Gαi3 (937 nM) or the non-phosphorylatable 3YF mutant and immunoblotted with custom rabbit polyclonal anti-pYpY-Gαi3 antibody. All immunoblots are representative of findings from at least 3 independent repeats.

### Phosphorylation within the interdomain cleft enhances Gαi activation

Of the three phosphosites, Y320 is on β6 strand facing more towards the solvent (Fig. 3B; ***SI Appendix*, Fig. S4a**), whereas Y154/155 are ‘buried’ and inaccessible to solvent in all presently available structures of Gαi (either active or inactive) in which the interdomain cleft is ‘closed’ (Fig. 3b-c). Because of their stoichiometric abundance in cells (~65 for pY154/pY155; Fig 3d) and enhanced phosphorylation *in vitro* under conditions that are permissive to nucleotide exchange (Fig 2d-e), we hypothesized that phosphorylation may require a more relaxed conformation, such as the nucleotide-free transition states with ‘opening’ of the interdomain cleft [as shown to coincide with nucleotide release that is triggered by the GPCRs (50–53) and as deduced by NMR studies with GIV-GEM (45)]. Homology modeling studies revealed that Y154/155 are still not accessible in the nucleotide-free ‘open’ state (***SI Appendix*, Fig. S4b**); however, the intricate network of hydrogen bonds that is facilitated by Y154/Y155 between the αE helix, αF helix, and Sw-I in the nucleotide-bound state is lost in the nucleotide-free state (***SI Appendix*, Fig. S4c**). Computational modeling predicted that phosphorylation at either tyrosine would also disrupt this network of hydrogen bonds (Fig. 4a) and destabilize the overall Gαi structure (Fig. 4b), suggesting that phosphorylation may induce hitherto unknown structural changes with important functional consequences. Phosphorylation did not appear to overtly impact nucleotide-binding, as determined by limited trypsin proteolysis assays (***SI Appendix*, Fig. S4d**). Despite multiple attempts, three different strategies to purify phosphorylated/phospho-mimicking Gαi3 failed: 1) size exclusion chromatography after *in vitro* phosphorylation; 2) replacement of Y with a pY-mimicking non-natural amino acid, p-carboxymethyl-L-phenylalanine (pCMF) (54) (see ***SI Appendix*, Fig. S4e**); and 3) replacement of Y154/155 with a Glu(E). These unsuccessful attempts indicate that the predicted degree of structural instability of these phosphotyrosines (Fig. 4b) may preclude protein purification altogether. Among the single-Y mutants, we were only able to generate Gαi3-Y154E with high purity and at low, but sufficient, amounts to proceed with use in functional assays; we also included the Gαi3-3YF mutant in these functional assays. After confirming that these WT and mutant proteins were capable of binding nucleotides and adopting an active conformation (***SI Appendix*, Fig. S3d and S4f**), we assessed their thermostability using a well-accepted approach, differential scanning fluorimetry (thermal shift assays) (55). The Y154E mutation impacted the stability of Gαi drastically; the melting temperature (Tm) of Gαi3-Y154E in the native state could not be determined (Fig. 4c) and was significantly lower in nucleotide-bound states compared to the WT protein [-10.75◻C Tm change for the GDP-bound state (Fig. 4d) and −7.5◻C Tm change for the GTPγS-bound state (***SI Appendix*, Fig. S4g**)]. Instability was also accompanied by increased rates of basal nucleotide exchange compared to Gαi3-WT (~15.6-fold increase; Fig. 4e-f). Notably, the Gαi3-3YF mutant, in which the Phe(F) cannot participate in H-bonds, exhibited thermal stability (Fig. 4c-d; ***SI Appendix*, Fig. S4g**) and basal nucleotide exchange rate (Fig. 4e-f) comparable to the WT protein. These findings suggest that the impact of phosphorylation on the stability of Gαi may not be attributed entirely to the loss of the H-bond network in the αE-αF region of the protein; the Y154E mutation (and by that token, phosphorylation induced changes) must alter other intramolecular interactions to account for the observed decrease in thermal stability and increase in activity.

**Fig. 4:**
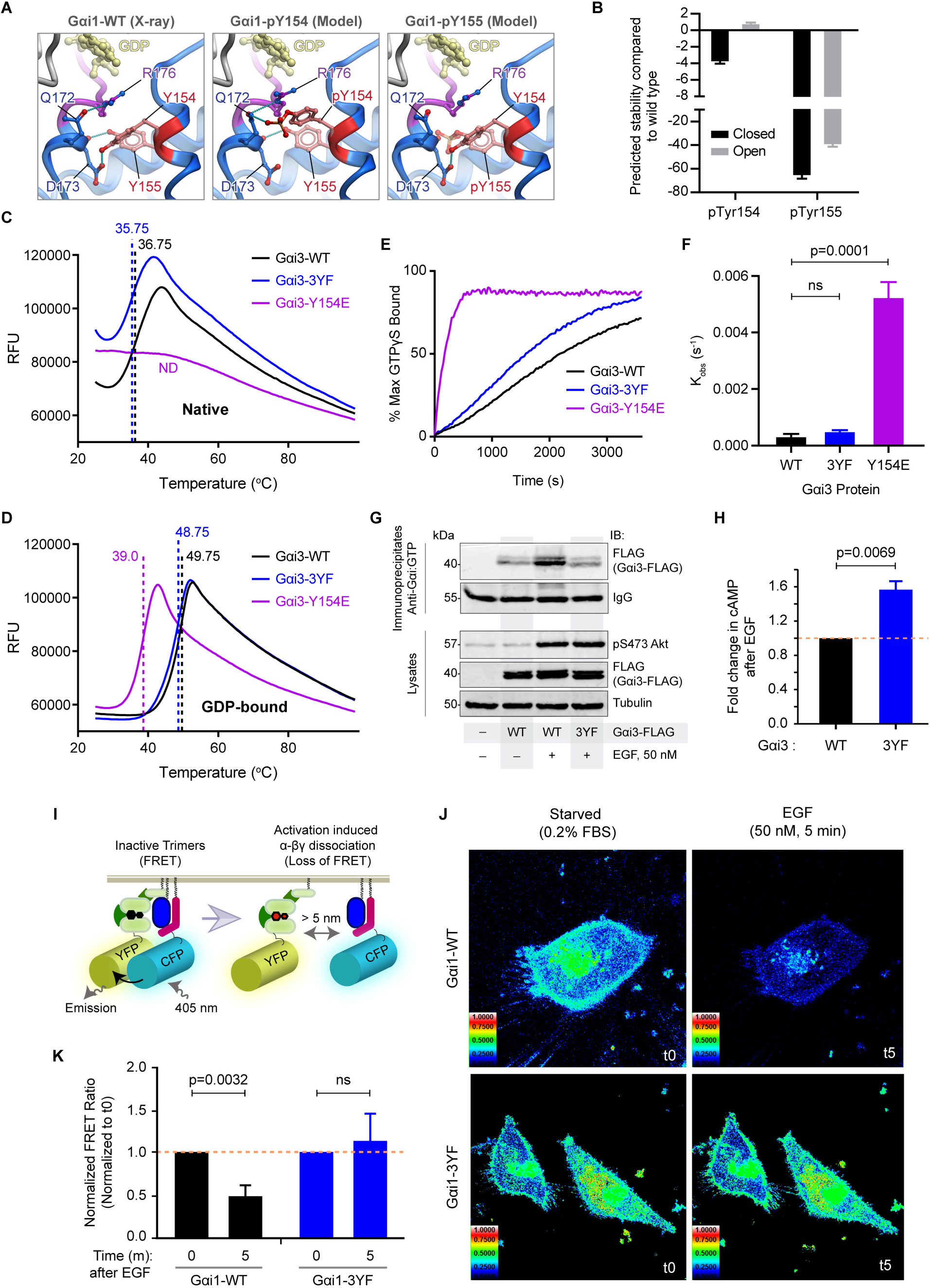
Phosphorylation of Gαi by RTKs is required for efficient Gαi activation and downstream signaling. **A**, Structures of Gαi1 highlighting the hydrogen bonding between Y154/Y155 and neighboring residues with and without being phosphorylated. **B**, Bar graph displaying computationally predicted structural stability of Gαi3-pY154 and Gαi3-pY155 in the ‘open’ and ‘closed’ states. **C-D**, WT, non-phosphorylatable (YF) and phosphomimicking (YE) mutant Gαi proteins were subjected to increasing temperatures in differential scanning fluorimetry (thermal shift) assay. Findings are displayed as a line graphs charting the average relative fluorescence units (RFU) of native (no excess GDP; **C**) and GDP-bound (1 mM GDP added; **D**) Gαi proteins. Measured melting temperature for each condition is indicated by the vertical dotted lines. **E-F**, GTPγS incorporation into WT and various mutant Gαi proteins listed in **C** was measured by intrinsic tryptophan fluorescence. Findings are displayed as a line graph (**E**) showing average nucleotide incorporation over time and a bar graph (**F**) showing the observed nucleotide incorporation rates (*k*_*obs*_, s^-1^). Error bars, ±S.E.M.; n = 3. **G**, Immunoprecipitation using the Gαi:GTP (active conformation specific) antibody of transfected FLAG-tagged Gαi3-WT or Gαi3-3YF from EGF-stimulated Cos-7 cells treated with PP2 inhibitor. Quantification shown in ***SI Appendix*, Fig. S5a**. **H**, Bar graph displaying fold change in cellular cAMP detected from HeLa Gαi3-WT or Gαi3-3YF EGF-stimulated cell lysates by radioimmunoassay. See also ***SI Appendix*, Fig. S5b**. Error bars, ±S.E.; n = 3. **I**, Schematic describing the mechanism of the FRET Gαi activity reporter. **J**, Representative FRET images of cells expressing Gαi1-WT (*top*) or Gαi1-3YF (*bottom*) activity reporters before and after EGF stimulation. FRET scale is shown inset. **K**, Bar graphs displaying quantification of FRET results from (**J**). Error bars, ±S.E.M.; n = 5-7 cells/experiment, from 4 independent experiments. All immunoblots are representative of findings from at least 3 independent repeats.

In the absence of any discernible functional defects *in vitro*, we used the non-phosphorylatable 3YF mutant in cell-based assays to assess activation of Gαi after EGF stimulation: 1) Direct assessment of Gαi•GTP (active) using a previously validated conformation-specific antibody (56) (Fig. 4g; ***SI Appendix*, Fig. S5a)**; 2) Indirect assessment of activation by monitoring ligand-stimulated suppression of cellular cAMP (57) by radioimmunoassay (RIA; Fig. 4h; ***SI Appendix*, Fig. S5b**); 3) A förster resonance energy transfer (FRET)-based approach where activation is indirectly monitored by the dissociation of fluorescent-tagged Gαi and Gβγ subunits with a resultant loss of FRET (58, 59) (Fig. 4i-k). Findings in all three cellular approaches concurred with our prior conclusions, i.e., phosphorylation increases Gαi activation. First, Gαi3-WT but not the Gαi3-3YF mutant was efficiently immunoprecipitated in EGF-stimulated cells by anti-Gαi•GTP antibody (Fig. 4g; ***SI Appendix*, Fig. S5a**). Second, RIA assays on Gαi-depleted HeLa cell lines stably expressing WT or the 3YF mutant Gαi3 (***SI Appendix*, Fig. S4h**) confirmed that Gαi3 suppresses cellular cAMP after EGF stimulation (***SI Appendix*, Fig. S5b**); however, the Gαi-3YF mutant was less efficient in doing so (~55% increase compared to Gαi3-WT; Fig. 4h). Finally, cells expressing Gαi-WT exhibited a robust loss of FRET signal (~50% decrease) and was efficiently activated in response to EGF stimulation, whereas cells expressing the Gαi-3YF mutant did not (Fig. 4j-k). These *in vitro* and cellular findings indicate that RTK phosphorylation of Gαi is required for efficient Gαi activation and signaling downstream of growth factors.

### Cancers harbor Gαi mutants that mimic constitutive activation of the RTK→Gαi pathway

A search of various catalogs of somatic mutations in cancers revealed that both Y154 and Y155 are mutated in different tumors (***SI Appendix*, Fig. S6a**). Computational modeling predicted each mutation to not only disrupt H-bond network within the inter-domain cleft but also destabilize the protein (Fig. 5a; ***SI Appendix*, Fig. S6b-c**). Consistent with these predictions and much like our limited success in expressing the pY-mimic pCMF or YE mutants, we were unsuccessful in generating all but one of these mutants (Fig. 5a; ***SI Appendix*, Fig. S6d**); the mutant that was predicted to be most stable among them all, Y154H, was purified at amounts that were sufficient to characterize in functional assays. Limited proteolysis assays confirmed that the Gαi3-Y154H mutant could bind nucleotides and adopt an active conformation (***SI Appendix*, Fig. S7a**). Thermal shift assays showed that the Y154H mutant was less stable compared to the WT protein in the native (−4◻C Tm change; Fig. 5c), and GTP-bound (−5◻C Tm change; ***SI Appendix*, Fig. S7b**) states. Nucleotide exchange assays revealed that instability of Y154H was accompanied by increased rates of basal nucleotide exchange (~5-fold higher than WT; Fig. 5d-e); this was also reflected in increased overall cycling in steady-state GTPase assays (***SI Appendix*, Fig. S7c**). To determine if the increase in exchange rate observed *in vitro* translates to constitutive (growth factor-independent) activation in cells, we carried out FRET-based Gαi activity reporter assays at steady state in the presence of reduced serum (2% FBS). Gαi3-Y154H had significantly higher activation, showing about an 80% reduction in FRET signal compared to Gαi3-WT and Gαi3-3YF (Fig. 5f-g; ***SI Appendix*, Fig. S7d**). In addition, no further change in FRET was seen in the Gαi3-Y154H mutant after EGF stimulation, indicating that ligand stimulation could not activate this mutant any further (***SI Appendix*, Fig. S7e-f**). These results demonstrate that cancer-associated somatic mutations at Y154, and perhaps also at Y155, regulate G protein activation in cells, and that such mutations may mediate ligand-independent constitutive activation of the pathway in tumors.

**Fig. 5:**
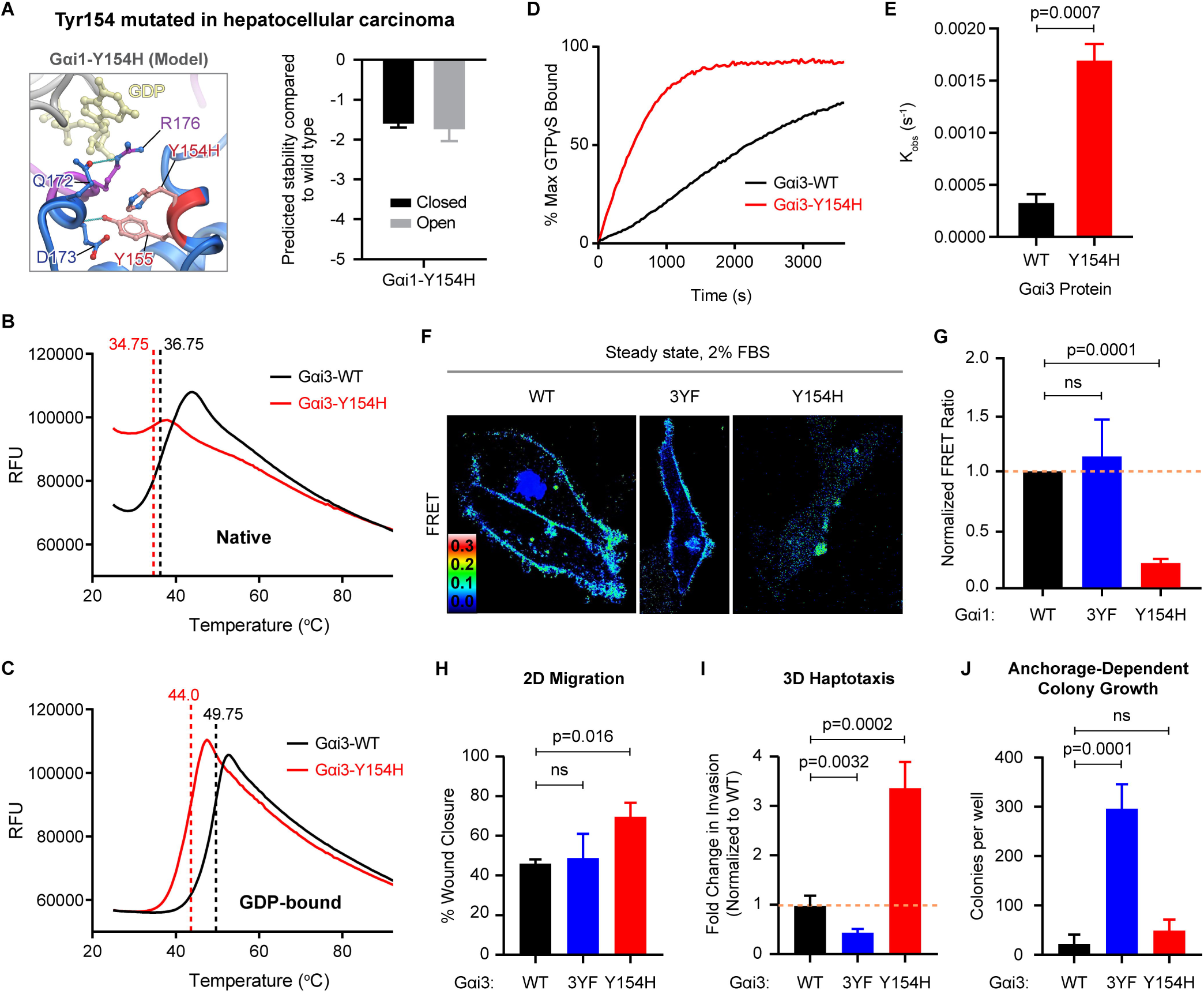
Cancer-associated mutations at Y154 in Gαi hyperactivates the G protein. **A**, (*Left*) Structural model of Gαi1-Y154H mutation; a comparison with **4A** highlights loss of H-bonds due to mutation. (*Right*) Bar graph displaying computationally predicted structural stability of Gαi1-Y154H in the open and closed states. **B-C**, WT and Y154H mutant Gαi proteins were subjected to increasing temperatures in differential scanning fluorimetry (thermal shift) assay. Findings are displayed as a line graph showing average relative fluorescence units (RFU) curves of native (no excess GDP; **B**) and GDP-bound (1 mM GDP added; **C**) Gαi proteins. Measured melting temperature for each condition is indicated by the vertical dotted lines. **D-E**, GTPγS incorporation into WT and Y154H mutant Gαi proteins measured by intrinsic tryptophan fluorescence. Findings are displayed as a line graph (**D**) showing average nucleotide incorporation over time and a bar graph (**E**) showing the observed nucleotide incorporation rates (*k*_*obs*_, s^-1^). Data shown is from three independent experiments. WT data shown in **B-E** is same as WT data shown in Figure 4C-F. **F**, Representative FRET images of cells expressing Gαi1-WT (*left*), Gαi1-3YF (Y154/155/320F; *middle*), and Gαi1-Y154H (*right*) activity reporters under steady state conditions with 2% FBS. Corresponding CFP and YFP images are shown in ***SI Appendix*, Fig. S7d**. **G**, Bar graphs displaying quantification of FRET results from (**F**). Error bars, ±S.E.M.; n = 11-19 cells/experiment, from 4 independent experiments. **H**, Bar graphs display % wound closure in 2D-scratch-wound assays performed using Gαi-depleted (by shRNA) HeLa cells stably expressing shRNA resistant rat Gαi-WT, 3YF and YH constructs. Error bars, ±S.E.M.; n = 3. Representative wound images are shown in ***SI App dix*, Fig. S7h**. **I**, Bar graphs display fold change in the number of cells that migrated across a 0-to-10% serum gradient in 3D-transwell assays performed using the same cell lines in as **H**. Error bars, ±S.E.M.; n = 3. Representative images of the porous transwell membrane are shown in ***SI Appendix*, Fig. S7j**. **J**, Bar graphs display the number of colonies/well in anchorage-dependent colony formation assays performed using the same cell lines in as **H**. Error bars, ±S.E.M.; n = 3. Representative colony formation assays are shown in ***SI Appendix*, Fig. S7k**.

### RTK-dependent phosphorylation of Gαi impacts cell behavior

Leveraging this hyperactive Y154H mutant as a tool, we asked how constitutive activation of this RTK→Gαi pathway may impact cell phenotype. Previous work showed GIV-dependent activation of Gαi enhances motility but inhibits proliferation, and thereby, orchestrates a migration-proliferation dichotomy (60, 61). Mechanistically, this dichotomy has been attributed to GIV’s ability to regulate the spatiotemporal aspects of EGFR signaling (from cell surface *vs*. endosomes) (60). In the presence of GIV, PM-based promotility pathways (PI3K-Akt) are enhanced but endosomal mitogenic signals (MAPK) are suppressed and hence, the cells migrate more than they proliferate; the reverse is true for cells without GIV or those without an intact GEM motif in GIV. Such dichotomy in 'Go or Grow' decision reflects a transition to invasive phenotypes that are triggered by stressors such as nutrient shortage within growing tumors (62–66). To assess how the phosphoevents identified here impact migration-proliferation dichotomy, we generated Gαi3-depleted HeLa cell lines stably expressing WT or Y154H mutant G protein (***SI Appendix*, Fig. S7g**) and assessed their ability to migrate in 2D-scratch wound and 3D-transwell assays and proliferate in anchorage-dependent colony growth assays. To mimic intratumoral conditions of limited growth factors, and consistent with all prior work assessing the functions of GIV’s GEF function downstream of RTKs (34, 39–41, 60), the 2D-scratch wound assays and colony formation assays were carried in the presence of 2% FBS while the 3D-transwell assay was conducted using a 0-to-10% serum gradient. Compared to the cells expressing Gαi3-WT, those expressing the Y154H mutant exhibited increased cell migration in the 2D-scratch wound assay (23.9% more area closure; Fig. 5h; ***SI Appendix*, Fig. S7h**) and in the 3D-transwell assay (~3x more; Fig. 5i; ***SI Appendix*, Fig. S7i-j**), but a similar amount of growth in colonies (Fig. 5j; ***SI Appendix*, Fig. S7k**). By contrast, in the same assays, cells expressing Gαi3-3YF migrated either to a similar extent (in 2D-scratch would assays; Fig. 5h; ***SI Appendix*, Fig. S7h**) or to a significantly lesser extent (~2x less in 3D-transwell assays compared to WT; Fig. 5i; ***SI Appendix*, Fig. S7i-j**) but showed enhanced growth in anchorage-dependent colony formation assays (~12x more colonies compared to WT; Fig. 5j; ***SI Appendix*, Fig. S7k**). Results indicate that in partially starved conditions constitutive phosphorylation (as mimicked by the Y154H mutant) favors migration, whereas a constitutive non-phosphorylated state (mimicked by the Y3F mutant) favors proliferation. From these we conclude that the RTK→Gαi pathway we report here is a key determinant of migration proliferation dichotomy and may serve as a key step within a cell’s decision-making process to ‘Go or Grow’. Enhanced signaling through this axis can drive a pro-metastatic tumor cell phenotype, either rarely *via* infrequently encountered Y154/155 mutations in Gαi or widely *via* the more frequently encountered elevated expression and hypersignaling via growth factor wRTKs.

## Discussion

The most important discovery we report here is defining the key steps of and the underlying molecular mechanisms that enable growth factor RTKs to trigger G protein signaling (Fig. 6a). Activation of inactive GDP-bound Gαiβγ trimers by RTKs requires first the scaffolding of ligand-activated RTKs (i.e., kinase) with monomeric Gαi (i.e., substrate) within RTK•GIV•Gαi ternary complexes; this step is facilitated by GIV. Because phosphorylation is significantly higher in cells than *in vitro*, and the latter is enhanced when kinase assays were carried out under conditions that favor nucleotide exchange, it is likely that the relatively buried Y154/Y155 in Gαi become more accessible during GIV-triggered allosteric conformational changes that ultimately lead to nucleotide exchange (44, 45). Findings also show that phosphorylation of Gαi on two tandem sites within the inter-domain cleft (Y154 and Y155), enhances the nucleotide exchange rate of Gαi, regulates cAMP and alters cellular phenotypes. Because RTKs signal primarily *via* tyrosine phosphorylation cascades, the evidence for tyrosine-based *transactivation* of G proteins we provide here represents a crosstalk between two most widely studied pathways in the field of signal transduction. By pinpointing the sequential protein-protein interactions that ultimately lead to the unique tyr-phosphorylation events on Gαi and interrogating the impact of those events on the stability and exchange rates of the GTPase, we have revealed the molecular/structural basis for this crosstalk. Using non-phosphorylatable and tumor-inspired constitutively phosphomimicking mutants we have obtained evidence for the existence of this paradigm in cells and charted the cellular consequences of such crosstalk in cancers.

**Fig. 6:**
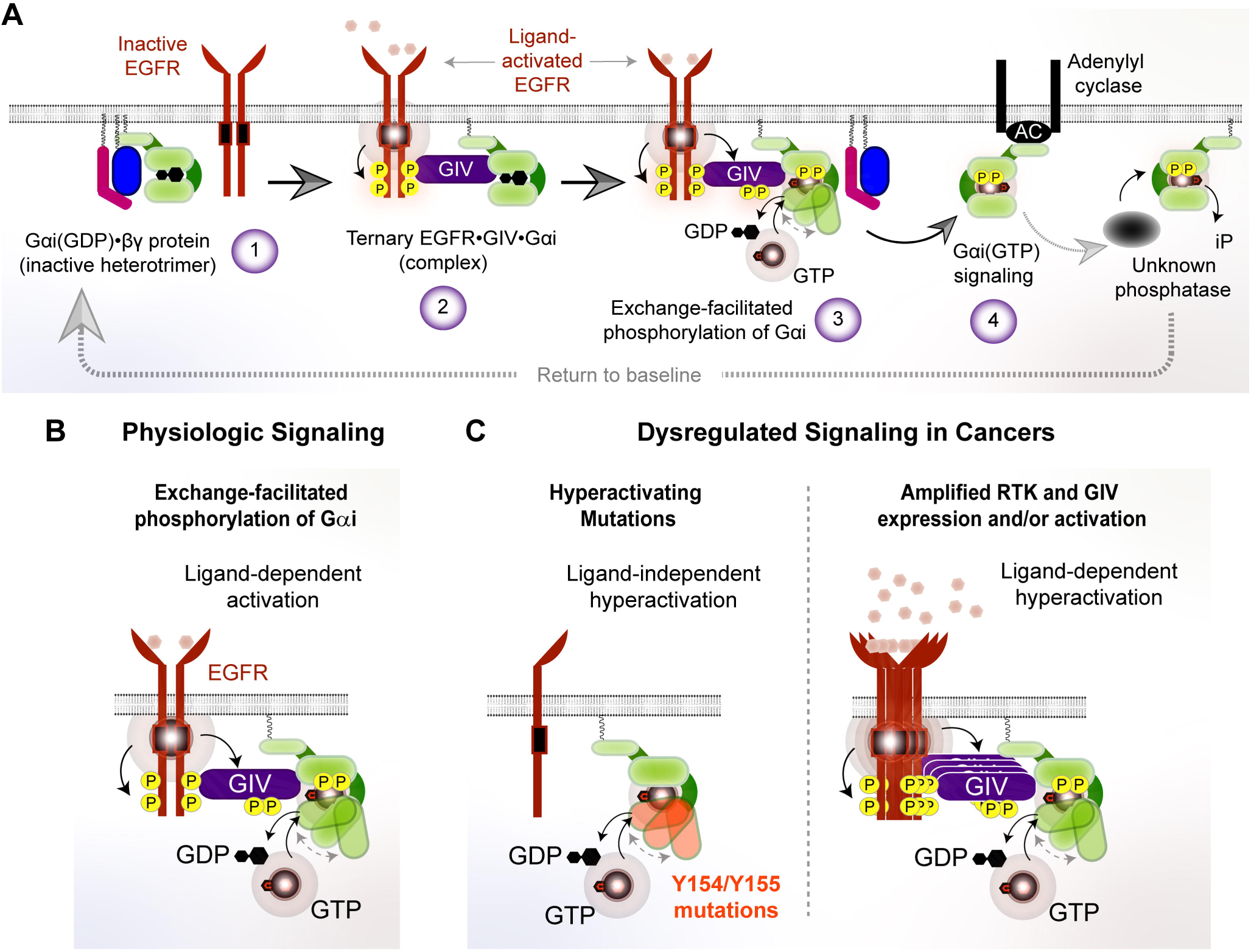
Summary of findings and their implications. **A**, Activation of inactive GDP-bound Gαiβγ (step 1) within this pathway requires its physical coupling to a ligand-activated RTK within an RTK•GIV•Gαi ternary complex (step 2), and subsequent phosphorylation of Gαi on two tandem sites within the inter-domain cleft (step 3). Phosphorylation enhances nucleotide exchange rate of Gαi, regulates cAMP (step 4) and alters cellular phenotypes. **B**, Physiologic growth factor-dependent activation of RTKs leads to phosphorylation and activation of Gαi and downstream signaling. **C**, Pathologic hyperactivation of this pathway can occur via activating mutations in Gαi (left), amplification of GIV or RTKs (right), or activating mutations in RTKs. Such hyperactivation can drive cancer progression and tumor cell phenotypes such as increased tumor cell migration.

The impact of these findings on the two large fields that they straddle is many fold: *First*, in the context of tyrosine signaling, all RTKs that bind GIV (e.g., InsR, PDGFR, VEGFR, etc.) may trigger this crosstalk provided they are activated *and* GIV is present in sufficient amount to scaffold the ligand-activated RTK (kinase) to the G protein (substrate). Such redundancy or versatility may represent a convergence point for signals, which is expected to confer phenotypic robustness. *Second*, in the context of G protein/GPCR signaling, that RTKs phosphorylate residues (Y154/155) that are buried within the inter-domain cleft implies that the residues must get exposed to solvent within a disordered/linear stretch that is accessible to the kinases (67, 68). That such exposure is not seen in any of the solved structures of Gαi in ‘open’ nucleotide-free conformation (all stabilized by Fab antibodies) implies that transition states of Gαi with a greater degree of unfolding of its helical domain (or at least of the αE helix) must exist that are yet to be discovered. The predicted and observed impact of phosphorylation at Y154/155 on protein stability (decrease) and the *in vitro* and in-cell evidence of their impact on basal nucleotide exchange rates (increase) suggests that phosphorylation within the interdomain cleft may affect its opening/closing. Although molecular dynamic simulation studies have shown that domain opening is insufficient for GDP release (69), such opening can affect overall protein stability. But how may GDP release be triggered? One possibility is that phosphorylation at Y154/155 may affect neighboring residues within the nucleotide-binding pocket, such as those in the αD-αE loop which includes the so-called ‘NDS’ motif. Alternatively, phosphorylation at Y154/155 may affect Sw-I *via* the αE-αF loop; Sw-I was recently identified as a key conduit in the allosteric path to nucleotide release when GIV binds Gαi, transmitting forces from the Sw-II to the hydrophobic core of the GTPase (44). Because both Y154 and Y155 face towards αF and Sw-I, particularly Y155, it is possible that destabilization of Sw-I could serve as a mechanism for pTyr-induced allosteric activation of the GTPase. If so, allosteric movements in Sw-I could be a shared mechanism for a potential synergy between GIV-dependent and pTyr-dependent G protein activation.

Because of its stoichiometrically determined minority status, the role of pY320 was not pursued here. Given its location on the β6-strand, which is within the GPCR-binding interface, it is tempting to speculate that pY320 may impact GPCR binding. Because Gαi-Y320 was independently identified recently as important for coupling to GPCRs (70), further investigation is warranted to see if RTK-dependent phosphorylation at Y320 builds upon the theme of cooperativity between RTKs and G proteins/GPCRs.

Finally, evidence presented here shows that the RTK→Gαi pathway may be hijacked in tumors to support sinister cell phenotypes. Related to tumor growth and metastasis, the unphosphorylatable Gαi-3YF mutant displayed enhanced anchorage-dependent colony formation but reduced haptotaxis compared to Gαi-WT, while the active Gαi-Y154H cancer mutant displayed enhanced 2D and 3D migration compared to the WT G protein. Although we demonstrate that ‘activating’ mutations in Gαi, such as the Gαi-Y154H that we characterized here, can turn on this crosstalk pathway, these mutations are likely to be rare events. However, the importance of this crosstalk in human cancer goes beyond just the Y154H mutant. Upregulation (by increasing copy #s) of GIV (71) or RTKs such as EGFR (72), or activating mutations of the latter (73), is a much more common event in tumors that could activate this TK-dependent phospho-activation of G proteins and result in resistance to EGFR-targeted therapies and poorer prognosis (Fig. 6b-c; ***SI Appendix*, Fig. S8-S9**). Given the well-known pharmacologic importance of the RTK and G/GPCR pathways (independent of each other), it is possible that the signaling interfaces that are uniquely assembled for the RTK→Gαi transactivation are of high value for tackling pathway crosstalk.

## Materials and Methods

Reagents and antibodies, plasmid constructs, mutagenesis and protein expression and purification, cell culture and immunoblotting, immunoprecipitation assays, *in silico* evaluation of effects of mutations and phosphoevents on Gαi stability, FRET, migration assays in 2D- and 3D migration assays and anchorage-dependent colon growth assays are detailed in *SI Appendix*, and briefly mentioned here.

### Cell culture, transfection, lysis, and quantitative immunoblotting

HeLa, Cos-7 and HEK293 cells were cultured according to American Type Collection guidelines. HeLa cell lines stably expressing Gαi3 wild-type (HeLa-Gαi3-WT), Gαi3-Y154/155/320F (HeLa-Gαi3-3YF), or Gαi3-Y154H were generated as described previously (41). GIV-depleted HeLa cell lines (by shRNA) stably expressing shRNA-resistant GIV-WT and GIV-FA mutants were previously generated and extensively validated through numerous studies interrogating the GIV•Gαi interface (34, 39, 41, 43, 60, 74).

For quantitative immunoblotting, infrared imaging with two-color detection and quantification were performed using a Li-Cor Odyssey imaging system. All Odyssey images were processed using Image J software (NIH) and assembled for presentation using Photoshop and Illustrator softwares (Adobe).

### In vitro kinase and in-cell phosphorylation assays

In vitro phosphorylation assays were carried out using purified His-Gαi3 wild-type or mutants (~1-5 µg/reaction) and commercially obtained recombinant kinases (50-100 ng/reaction). The reactions were started by addition of 1 mM of ATP and carried out at 25ºC in 50 µl of kinase buffer [60 mM Hepes (pH 7.5), 5 mM MgCl_2_, 5 mM MnCl_2_, 3 μM Na_3_OV_4_] for 60 min. For in vivo phosphorylation assays on Gαi3, Cos-7 cells were transfected with Gαi3-FLAG wild-type or mutants and serum-starved for 16 h (0 % FBS) prior to stimulation with EGF (50 nM, 5 min) or insulin (100 nM, 5 min) in the presence or absence of PP2 (10 μM, added 1 hr prior to stimulation. Reactions were stopped using PBS that was chilled to 4 ºC and supplemented with 200 μM sodium orthovanadate, and immediately scraped and lysed for immunoprecipitation followed by immublotting.

### Linear-ion-trap Mass Spectrometry

To determine *in vivo* phosphorylation states of the FLAG-Gαi3 we used the QTRAP 5500 in the selected reaction monitoring (SRM) mode to scan for all possible phospho-forms of this protein. For this purpose, SRM methods were developed for all possible tryptic peptides in phosphorylated and non-phosphorylated states [EYQLNDSASY^154^Y^155^LNDLDR and EVY^320^THFTCATDTK]. The ABSCIEX SRM Pilot™ software was used for SRM method development. Ultimately a method with 210 SRM transitions states was developed for phosphorylated and non-phosphorylated tryptic peptides of Gαi3 [**Dataset S1**]. In most cases there were at least 2 transitional states used for a given peptide mass. A total of 13 unique phosphorylation sites in the Gαi3 protein were detected by the QTRAP 5500, of which, 3 were tyrosines; all three tyrosines were detected also in His-Gαi3 protein that was phosphorylated *in vitro* by recombinant EGFR [**Dataset S2]**. Because samples were not subjected to phosphoenrichment prior to Mass Spectrometry analyses, stoichiometry of any phosphoevent was calculated based on the phosphorylated over total peptides of any given sequence.

To explore the possibility of the presence of other phosphorylation sites in Gαi3 protein, we used another 10 μL of the same tryptic sample used in the previous SRM experiment, to run the QTRAP 5500 mass spectrometer in the “precursor ion scanning mode” either for an ion at m/z 79 in negative ion mode for serine and threonine phosphorylation, or an ion at m/z 216.043 for tyrosine phosphorylation in the positive ion mode. Once the precursor ions are detected, the instrument switches to positive ion trap scanning mode to isolate the parent ions and to carry out MS2 analysis on these ions. The collected MS2 spectra were analyzed using the ProteinPilot^®^ search engine to identify the matching protein sequence from a database.

### Limited trypsin proteolysis assays

His-Gαi3 wild-type or His-Gαi3 mutants (0.5 mg/ml) were incubated for 120 min at 30 °C in the presence of GDP (30 µM) or GDP-AlF_4_^-^ (30 µM GDP, 30 µM AlCl_3_, 10 mM NaF). After incubation, samples were first in vitro phosphorylated by EGFR or directly treated with trypsin (final concentration, 12.5 µg/ml) and incubated for an additional 10 min at 30 °C. Reactions were stopped by adding SDS-PAGE sample buffer and boiling. Proteins were resolved by SDS-PAGE and stained with Coomassie Blue and/or immunoblotted with specific antibodies.

### In silico evaluation of effects of mutations and phosphoevents on Gαi stability

The stability changes in Gαi following Tyr phosphorylation or mutation were predicted by calculating the change in free energy compared to WT Gαi in ICM (Molsoft LLC), using either open (PDB: 6cmo (53), 6ot0 (52)), or closed (PDB: 1bof (75), 1gdd (76), 1gfi (77), 1gia (77), 1git (78), 1svk (79), 2hlb (80)) Gαi conformations. Briefly, Tyr residues were either phosphorylated (pTyr) or mutated to the respective residues (His, Ser or Asp) *in silico*, after which the mutated- and neighboring (within 6 Angstrom) residue side chains were sampled by Biased Probability Monte Carlo in internal coordinates (81). The free energy of folding (ΔG) for either WT or mutant protein in relation to the unfolded state of the same protein was approximated as a sum of empirical residue-specific energies previously optimized against a large experimental dataset (82). The difference ΔΔG = ΔG_WT_-ΔG_mut_ was used to estimate the destabilizing effect of the mutation or the phosphorylation event, with positive and negative numbers indicative of stabilization or destabilization, respectively. For pTyr substitutions, an earlier version of the algorithm was used where the resulting stability scores are measured on a different scale as compared to the newer version used for mutations; as a result, the number in Figure 4b are comparable to each other but not to the numbers in Figures 5a and others.

### Differential scanning fluorimetry (thermal shift assays)

His-Gαi3 (5 μM) was taken in their native state (as purified) or nucleotide loaded by incubating it for 150◻min at 30◻°C in buffer (20◻mM HEPES, pH 8, 100◻mM NaCl, 1◻mM EDTA, 10◻mM MgCl2, and 1◻mM DTT) supplemented with 1 mM GDP or 40 μM GTPγS. After loading, 45 μL of 5 μM His-Gαi3 was pipetted into PCR tubes (in triplicates) and 5 μL 200X SYPRO Orange solution freshly made in the same buffer from 5000X stock (Life Technologies S-6650) was added to the protein. A buffer + dye only (no protein) control was also included. Thermal shift assays were run on an Applied Biosystems StepOnePlus Real-Time PCR machine. Mixed protein and dye samples were subjected to increasing temperatures from 25 to 95°C in half degree increments, holding each temperature for 30 sec and measuring SYPRO fluorescence (using filter 3 for TAMRA™ and NED™ dyes) at each temperature. Melting temperatures were defined as the temperature at which the maximum value for the derivative of signal fluorescence (dF/dt) is achieved (GraphPad Prism v.7).

### GTPγS incorporation assays

A volume of 72 μL of His-Gαi3 at 1 μM in reaction buffer (20◻mM HEPES pH 8, 100◻mM NaCl, 1◻mM EDTA, 10◻mM MgCl_2_, and 1◻mM DTT) was transferred to a pre-warmed 384-well black flat-bottom plate (in triplicates). The reaction was initiated by injecting 8 μL of 1 mM GTPγS (Abcam, Cambridge, MA) in each well for a final reaction volume of 80 μL and final concentrations of 100 nM Gαi3, 100 µM GTPγS. Reactions were carried out at 30°C. GTPγS incorporation into Gαi3 was quantified by direct tryptophan fluorescence (ex = 280; em = 350), using a microplate fluorescence reader (TECAN Spark 20M). Fluorescence was measured every 30 sec starting immediately after injection of GTPγS. Raw fluorescence was plotted over time and observed rates (k_obs_) were determined by fitting a one-phase association curve to the data (GraphPad Prism v.7).

### Measurement of cAMP by radioimmunoassay

HeLa cells stably expressing rGαi3-WT or rGαi3-3YF were depleted of endogenous Gαi3 using siRNA, serum starved (0.2 % FBS, 16 h) and incubated with isobutylmethylxanthine (IBMX, 200 μM, 20 min) followed by EGF (50 nM, 10 min) and Forskolin (10 μM, 10 min). Reactions were terminated by aspiration of media and addition of 150 μl of ice-cold TCA 7.5% (w/v). cAMP content in TCA extracts was determined by radioimmunoassay (RIA) (83) and normalized to protein [(determined using a dyebinding protein assay (Bio-Rad)]. Data is expressed as fold change over Forskolin stimulation.

### F◻rster Resonance Energy Transfer (FRET) studies

Intramolecular FRET was detected by sensitized emission using the three-cube method were performed exactly as previously reported by Midde et al (33) and is detailed in *SI*.

### 2D scratch wound migration assay

Scratch-wound assays were done as described previously (38) and detailed in *SI*.

### 3D Transwell migration assay

These assays were done as described previously (84) and detailed in *SI*.

### Anchorage-dependent colony formation assay

Anchorage-dependent growth was monitored as described previously (85) and detailed in *SI*.

### Statistical analysis

Each experiment presented in the figures is representative of at least three independent repeats (with at least two technical repeats for each condition within each repeat). Statistical significance between the differences of means was calculated using multiple comparisons in one-way nonparametric ANOVA. All statistics and graphical data presented were prepared using GraphPad Prism v.7. All error bars are standard deviation.

## Acknowledgements

We thank Bridgett Simmons (AB SCIEX, CANADA) for technical assistance with mass spectrometry experiments and Yelena Pavlova and Nina Sun for technical assistance with cloning and mutagenesis of constructs. This paper was supported by the NIH (CA238042, AI141630, CA100768 and CA160911 to P.G). N.A.K was supported by a NIH predoctoral fellowship (F31 CA206426), and T32 training grants T32CA067754 and T32DK007202. M.G-M. was supported by the NIH (GM136132 and GM130120). I.L.-S. was supported by a fellowship from the American Heart Association (AHA 14POST20050025). I.K was supported by the NIH (AI118985 and R01 GM117424). T.N. is supported by a NHMRC C. J. Martin Early Career Fellowship 1145746.

