## Supplementary Online Materials for "Receptor tyrosine kinases activate heterotrimeric G proteins via phosphorylation within the interdomain cleft of Gαi"

**Supplementary Methods**
**Supplementary Figures** = 9
**Supplementary Datasets** = 2

### Supplementary Detailed Methods

##### *Reagents and antibodies*

Unless otherwise indicated, all reagents were of analytical grade and obtained from Sigma-Aldrich. Cell culture media were purchased from Invitrogen. EGF and insulin were obtained from Invitrogen and Novagen, respectively. Recombinant EGFR, PDGFR, VEGFR, InsR and Src were purchased from Cell Signaling Technology. PP2 was obtained from Calbiochem. Silencer Negative Control scrambled (Scr) siRNA and Gαi3 siRNA were purchased from Ambion and Santa Cruz Biotechnology, respectively. Antibodies against GIV that were used in this work include affinity-purified anti-GIV coiled-coil IgG and anti-GIV-CT (Santa Cruz Biotechnology). Anti-Gαi3:GTP (6-F12) was a gift from Dr. Graeme Milligan. Rabbit polyclonal antibodies used in this work were Gαi3 (C-10) and pan-Gβ (M-14) (Santa Cruz Biotechnology) and phospho-AKT Ser 473 (Cell Signaling). The rabbit anti-Gαi-pY154/pY155 (pYpY-Gαi) antibody was custom made ($21^{st}$ Century) for use in these studies. Mouse mAbs against pTyr (BD Biosciences), hexahistidine, FLAG and α-tubulin (Sigma) were obtained commercially. For immunoprecipitation assays, control mouse and rabbit IgGs were purchased from Bio-Rad and Sigma, respectively.

##### *Plasmid constructs, mutagenesis and protein expression*

Cloning of Gαi1/2/3, Gαs, Gαo and GIV-CT into pGEX-4T-1 or pET28b and rat Gαi3 and GIV into p3XFLAG-CMV$^{TM}$-14 were described previously (1). GST–TrkA-CT (aa 448-552) was a gift from Dr M.G. Farquhar and was used previously (1, 2). Mutants of rat Gαi3 (Y155F, Y320F, Y155/320F and Y155/155/320F) were generated using QuickChange II (Stratagene) and specific primers (sequence available upon request) following the manufacturer´s instructions and Gαi3-W258F-FLAG was described before (3). All constructs were checked by DNA sequencing.

***Expression and purification of His- and GST-tagged proteins***

His-tagged constructs were expressed and purified from *Escherichia coli* strain BL21 (DE3; Invitrogen) as described previously. Briefly, 1L of cells were grown in 2 L flasks at 37°C until OD reached 0.8-1.0, then induced overnight at 25°C with 1 mM IPTG. A bacterial pellet from 1 L of culture was resuspended in 10 ml of His lysis buffer (50 mM $NaH_2PO_4$ pH 7.4, 300 mM NaCl) supplemented with 2x Protease Inhibitors (Roche Life Science). Cell lysates were sonicated (4 × 20 s, 1 min between cycles) and then centrifuged at 12,000 × *g* at 4°C for 20 min. Solubilized proteins were affinity purified on His (cobalt or nickel) beads (GE Healthcare) by incubation for 4 hours at 4°C. Beads were washed 3 x with 50mM Tris pH 8 and then eluted with His elution buffer (50 mM $NaH_2PO_4$ pH 7.4, 300 mM NaCl, 300 mM imidazole). Eluted proteins were dialyzed overnight at 4°C against phosphate-buffered saline (PBS), and stored at −80°C in aliquots. His-tagged Gαi3 proteins were stored in storage buffer containing 20 mM Tris-HCl pH 7.4, 20 mM NaCl, 1 mM $MgCl_2$, and 5% glycerol).

GST-tagged constructs were expressed and purified from *Escherichia coli* strain BL21 (DE3; Invitrogen) as described previously. Briefly, 1L of cells were grown in 2 L flasks at 37°C until OD reached 0.8-1.0, then induced overnight at 25°C with 1 mM IPTG. A bacterial pellet from 1 L of culture was resuspended in 10 ml of GST-lysis buffer (25 mM Tris-HCl, pH 7.5, 20 mM NaCl, 1 mM EDTA, 20% [vol/vol] glycerol, 1% [vol/vol] Triton X-100, 2× protease inhibitor cocktail [Complete EDTA-free; Roche Diagnostics]). Cell lysates were sonicated (4 × 20 s, 1 min between cycles) and then centrifuged at 12,000 × *g* at 4°C for 20 min. Solubilized proteins were affinity purified on glutathione-Sepharose 4B beads (GE Healthcare) by incubation for 4 hours at 4°C. Beads were washed 3 x with 50mM Tris pH 8 and then eluted with GST elution buffer (50 mM Tris pH 8, 10 mM reduced glutathione). Eluted proteins were dialyzed overnight at 4°C against phosphate-buffered saline (PBS), and stored at −80°C in aliquots. GST-tagged Gαi3 proteins were stored in storage buffer containing 20 mM Tris-HCl pH 7.4, 20 mM NaCl, 1 mM $MgCl_2$, and 5% glycerol).

***Cell culture, transfection, lysis, and quantitative immunoblotting***

HeLa, Cos-7 and HEK293 cells were cultured according to American Type Collection guidelines. Cells were transfected using Genejuice (Novagen) or polyethylenimine for DNA plasmids and Oligofectamine (Invitrogen) for siRNA oligos following the manufacturers' protocols. HeLa cell lines stably expressing Gαi3 wild-type (HeLa-Gαi3-WT), Gαi3-Y154/155/320F (HeLa-Gαi3-3YF), or Gαi3-Y154H were generated as described previously (1). These cell lines were maintained in the presence of G418 (500 µg/ml). Lysates for immunoprecipitation assays were prepared by resuspending cells in lysis buffer [20 mM HEPES, pH 7.2, 5 mM Mg-acetate, 125 mM K-acetate, 0.4% Triton X-100, 1 mM DTT, supplemented with sodium orthovanadate (500 µM), phosphatase (Sigma) and protease (Roche) inhibitor cocktails], after which they were passed through a 28G needle at 4 °C, and cleared (10,000 x g for 10 min) before use in subsequent experiments. GIV-depleted HeLa cell lines (by shRNA) stably expressing shRNA-resistant GIV-WT and GIV-FA mutants were previously generated and extensively validated through numerous studies interrogating the GIV•Gαi interface (1, 4-8).

For immunoblotting, protein samples were separated by SDS-PAGE and transferred to PVDF membranes (Millipore). Membranes were blocked with PBS supplemented with 5% nonfat milk (or with 5% BSA when probing for phosphorylated proteins) before incubation with primary antibodies. Infrared imaging with two-color detection and quantification were performed using a Li-Cor Odyssey imaging system. All Odyssey images were processed using Image J software (NIH) and assembled for presentation using Photoshop and Illustrator softwares (Adobe).

***In vitro kinase and in-cell phosphorylation assays***

In vitro phosphorylation assays were carried out using purified His-Gαi3 wild-type or mutants (~1-5 µg/reaction) and commercially obtained recombinant kinases (50-100 ng/reaction). The reactions were started by addition of 1 mM of ATP and carried out at 25ºC in 50 µl of kinase buffer [60 mM Hepes (pH 7.5), 5 mM $MgCl_2$, 5 mM $MnCl_2$, 3 µM $Na_3OV_4$] for 60 min. Phosphorylated His-tagged proteins were detected by immunoblotting with mouse pTyr antibody and the total amount of proteins used in the assay were visualized by Ponceau S staining. For in vivo phosphorylation assays on Gαi3, Cos-7 cells were transfected with Gαi3-FLAG wild-type or mutants and serum-starved for 16 h (0 % FBS) prior to stimulation with EGF (50 nM, 5 min) or insulin (100 nM, 5 min) in the presence or absence of PP2 (10 µM, added 1 hr prior to stimulation. Reactions were stopped using PBS that was chilled to 4 ºC and supplemented with 200 μM sodium orthovanadate, and immediately scraped and lysed for immunoprecipitation followed by immublotting.

***Immunoprecipitation***

Cos-7 or HeLa cell lysates (~1–2 mg of protein) were incubated 3 h at 4 °C with 2 µg of antibody (FLAG, Gαi3 or Gαi3:GTP, depending on the experiment) followed by incubation with protein G or A-agarose beads at 4 °C for an additional 60 min. Beads were washed (x4) with 1 ml of wash buffer (4.3 mM $Na_2HPO_4$, .4 mM $KH_2PO_4$, pH 7.4, 137 mM NaCl, 2.7 mM KCl, 0.1% (v/v) Tween 20, 10 mM $MgCl_2$, 5 mM EDTA, 2 mM DTT), and the bound immune complexes were eluted by boiling in SDS sample buffer.

***Linear-ion-trap Mass Spectrometry***

To determine *in vivo* phosphorylation states of the FLAG-Gαi3 we used the QTRAP 5500 in the selected reaction monitoring (SRM) mode to scan for all possible phospho-forms of this protein. For this purpose, SRM methods were developed for all possible tryptic peptides in phosphorylated and non-phosphorylated states [EYQLNDSASY$^{154}$Y$^{155}$LNDLDR and EVY$^{320}$THFTCATDTK]. The ABSCIEX SRM Pilot™ software was used for SRM method development. Ultimately a method with 210 SRM transitions states was developed for phosphorylated and non-phosphorylated tryptic peptides of Gαi3 [**Supplementary Dataset S1**]. In most cases there were at least 2 transitional states used for a given peptide mass. A total of 13 unique phosphorylation sites in the Gαi3 protein were detected by the QTRAP 5500, of which, 3 were tyrosines; all three tyrosines were detected also in His-Gαi3 protein that was phosphorylated *in vitro* by recombinant EGFR [**Supplementary Dataset S2]**. Because samples were not subjected to phosphoenrichment prior to Mass Spectrometry analyses, stoichiometry of any phosphoevent was calculated based on the phosphorylated over total peptides of any given sequence.

To explore the possibility of the presence of other phosphorylation sites in Gαi3 protein, we used another 10 µL of the same tryptic sample used in the previous SRM experiment, to run the QTRAP 5500 mass spectrometer in the "precursor ion scanning mode" either for an ion at m/z 79 in negative ion mode for serine and threonine phosphorylation, or an ion at m/z 216.043 for tyrosine phosphorylation in the positive ion mode. Once the precursor ions are detected, the instrument switches to positive ion trap scanning mode to isolate the parent ions and to carry out MS2 analysis on these ions. The collected MS2 spectra were analyzed using the ProteinPilot® search engine to identify the matching protein sequence from a database.

***Limited trypsin proteolysis assays***

His-Gαi3 wild-type or His-Gαi3 mutants (0.5 mg/ml) were incubated for 120 min at 30 °C in the presence of GDP (30 μM) or GDP-$AlF_4^-$ (30 μM GDP, 30 μM $AlCl_3$, 10 mM NaF). After incubation, samples were first in vitro phosphorylated by EGFR or directly treated with trypsin (final concentration, 12.5 μg/ml) and incubated for an additional 10 min at 30 °C. Reactions were stopped by adding SDS-PAGE sample buffer and boiling. Proteins were resolved by SDS-PAGE and stained with Coomassie Blue and/or immunoblotted with specific antibodies.

***In silico evaluation of effects of mutations and phosphoevents on Gαi stability***

The stability changes in Gαi following Tyr phosphorylation or mutation were predicted by calculating the change in free energy compared to WT Gαi in ICM (Molsoft LLC), using either open (PDB: 6cmo (9), 6ot0 (10)), or closed (PDB: 1bof (11), 1gdd (12), 1gfi (13), 1gia (13), 1git (14), 1svk (15), 2hlb (16)) Gαi conformations. Briefly, Tyr residues were either phosphorylated (pTyr) or mutated to the respective residues (His, Ser or Asp) *in silico*, after which the mutated- and neighboring (within 6 Angstrom) residue side chains were sampled by Biased Probability Monte Carlo in internal coordinates (17). The free energy of folding (ΔG) for either WT or mutant protein in relation to the unfolded state of the same protein was approximated as a sum of empirical residue-specific energies previously optimized against a large experimental dataset (18). The difference $\Delta\Delta G = \Delta G_{WT} - \Delta G_{mut}$ was used to estimate the destabilizing effect of the mutation or the phosphorylation event, with positive and negative numbers indicative of stabilization or destabilization, respectively. For pTyr substitutions, an earlier version of the algorithm was used where the resulting stability scores are measured on a different scale as compared to the newer version used for mutations; as a result, the number in Figure 4b are comparable to each other but not to the numbers in Figures 5a and others.

***Differential scanning fluorimetry (thermal shift assays)***

His-Gαi3 (5 μM) was taken in their native state (as purified) or nucleotide loaded by incubating it for 150 min at 30 °C in buffer (20 mM HEPES, pH 8, 100 mM NaCl, 1 mM EDTA, 10 mM MgCl2, and 1 mM DTT) supplemented with 1 mM GDP or 40 μM GTPγS. After loading, 45 μL of 5 μM His-Gαi3 was pipetted into PCR tubes (in triplicates) and 5 μL 200X SYPRO Orange solution freshly made in the same buffer from 5000X stock (Life Technologies S-6650) was added to the protein. A buffer + dye only (no protein) control was also included. Thermal shift assays were run on an Applied Biosystems StepOnePlus Real-Time PCR machine. Mixed protein and dye samples were subjected to increasing temperatures from 25 to 95°C in half degree increments, holding each temperature for 30 sec and measuring SYPRO fluorescence (using filter 3 for TAMRA™ and NED™ dyes) at each temperature. Melting temperatures were defined as the temperature at which the maximum value for the derivative of signal fluorescence (dF/dt) is achieved (GraphPad Prism v.7).

***GTPγS incorporation assays***

A volume of 72 μL of His-Gαi3 at 1 μM in reaction buffer (20 mM HEPES pH 8, 100 mM NaCl, 1 mM EDTA, 10 mM $MgCl_2$, and 1 mM DTT) was transferred to a pre-warmed 384-well black flat-bottom plate (in triplicates). The reaction was initiated by injecting 8 μL of 1 mM GTPγS (Abcam, Cambridge, MA) in each well for a final reaction volume of 80 μL and final concentrations of 100 nM Gαi3, 100 μM GTPγS. Reactions were carried out at 30°C. GTPγS incorporation into Gαi3 was quantified by direct tryptophan fluorescence (ex = 280; em = 350), using a microplate fluorescence reader (TECAN Spark 20M). Fluorescence was measured every 30 sec starting immediately after injection of GTPγS. Raw fluorescence was plotted over time and observed rates ($k_{obs}$) were determined by fitting a one-phase association curve to the data (GraphPad Prism v.7).

***Steady-state GTPase Assay***

This assay was performed as described previously (1, 3, 19). Briefly, His-Gαi3 (100 nM) was diluted in in assay buffer [20 mM Na-HEPES, pH 8, 100 mM NaCl, 1 mM EDTA, 25 mM $MgCl_2$, 1 mM DTT, 0.05% (w:v) $C_{12}E_{10}$] and GTPase reactions initiated at 30°C by adding an equal volume of assay buffer containing 1 μM [γ-$^{32}$P]GTP (~50 c.p.m/ fmol). Duplicate aliquots (25μl) were removed at the times indicated in the figures or figure legends and the reaction was stopped by the addition of 975 μl ice cold 5% (wt:vol) activated charcoal in 20 mM $H_3PO_4$ (pH 3). Samples were centrifuged for 10 min at 10,000g, and 500 μl of the resultant supernatants were scintillation counted to determine the amount of [$^{32}$P]Pi released. Results are presented as raw counts per minute (c.p.m.).

***Measurement of cAMP by radioimmunoassay***

HeLa cells stably expressing rGαi3-WT or rGαi3-3YF were depleted of endogenous Gαi3 using siRNA, serum starved (0.2 % FBS, 16 h) and incubated with isobutylmethylxanthine (IBMX, 200 μM, 20 min) followed by EGF (50 nM, 10 min) and Forskolin (10 μM, 10 min). Reactions were terminated by aspiration of media and addition of 150 μl of ice-cold TCA 7.5% (w/v). cAMP content in TCA extracts was determined by radioimmunoassay (RIA) (20) and normalized to protein [(determined using a dyebinding protein assay (Bio-Rad)]. Data is expressed as fold change over Forskolin stimulation.

***Förster Resonance Energy Transfer (FRET) studies***

Intramolecular FRET was detected by sensitized emission using the three-cube method were performed as previously reported by Midde et al (21). All fluorescence microscopy assays were performed on single cells in mesoscopic regime to avoid inhomogeneities from samples as shown previously by Midde et al. (21, 22). Briefly, cells were sparsely split into sterile 35 mm MatTek glass bottom dishes and transfected with 1 μg of indicated constructs. To optimize the signal-to-noise ratio in FRET imaging, various expression levels of the transfected FRET probes were tested. However, to minimize complexities arising from molecular crowding, FRET probes were overexpressed by ∼1.5- to twofold compared with the endogenous proteins. Because the stoichiometry of FRET probes has a significant impact on FRET efficiency, cells that expressed equimolar amounts of donor and acceptor probes (as determined by computing the intensity of the fluorescence signal by a photon-counting histogram) were chosen selectively for FRET analyses. An Olympus IX81 FV1000 inverted confocal laser scanning microscope was used for live cell FRET imaging (UCSD-Neuroscience core facility). The microscope is stabilized on a vibration proof platform, caged in temperature controlled (37°C) and CO2 (5%) supplemented chamber. A PlanApo 60x 1.40 N.A. oil immersed objective designed to minimize chromatic aberration and enhance resolution for 405-605 nm imaging was used. Olympus Fluoview inbuilt software was used for data acquisition. A 515 nm Argon-ion laser was used to excite EYFP and a 405 nm laser diode was used to excite ECFP as detailed by Claire Brown's group (23). Spectral bleed-through coefficients were determined through FRET-imaging of donor-only and acceptor-only samples (i.e. cells expressing a single donor or acceptor FP). Enhanced CFP emission was collected from 425-500 nm and EYFP emission was collected through 535-600 nm and passed through a 50 nm confocal pinhole before being sent to a photomultiplier tube to reject out of plane focused light. Every field of view (FOV) is imaged sequentially through ECFPex/ECFPem, ECFPex/EYFPem and EYFPex/EYFPem (3 excitation and emission combinations) and saved as donor, FRET and acceptor image files through an inbuilt wizard. To obtain the FRET images and efficiency of energy transfer values a RiFRET plugin in Image J software was used (24). Prior to FRET calculations, all images were first corrected for uneven illumination, registered, and background-subtracted. Manual and automatic registration of each individual channel in ImageJ was critical to correct for motion artifacts associated with live cell imaging. Controls were performed in which images were obtained in different orders. The order in which images were obtained had no effect. FRET images were obtained by pixel-by-pixel ratiometric intensity method and efficiency of transfer was calculated by the ratio of intensity in transfer channel to the quenched (corrected) intensity in the donor channel. The following corrections were applied to all FOVs imaged: For cross-talk correction, cells transfected with CFP or YFP alone were imaged under all three previously mentioned excitation and emission combinations. FRET efficiency was quantified from 3-4 Regions of Interests (ROI) per cell drawn exclusively along the P.M. Because expression of FRET probes may have a significant impact on FRET efficiency, cells that expressed similar amounts of probes, as determined by computing the fluorescence signal/intensity by a photon counting histogram were selectively chosen for FRET analyses. Furthermore, untransfected cells and a field of view with-out cells were imaged to correct for background, autofluorescence and light scattering. To avoid artifacts of photobleaching, Oxyfluor (www.oxyrase.com) was used to minimize the formation of reactive oxygen species.

##### *2D scratch wound migration assay*

Scratch-wound assays were done as described previously (25). Briefly, monolayer cultures (100% confluent) of HeLa cells expressing WT, 3YF, or Y154H Gαi3 were scratch-wounded using a 10-µl pipette tip and incubated in 2% FBS media. The cells were subsequently monitored by phase-contrast microscopy over the next 30 hr. To quantify cell migration (expressed as percent of wound closure), images were analyzed using ImageJ software to calculate the difference between the wound area at 0 hrs and that at 20 hrs divided by the area at 0 hrs × 100.

##### *3D Transwell invasion assay*

Cell invasion was assessed using Transwell® Polycarbonate (PC) translucent (Corning, NY, USA) inserts with 8-µm pores in 24-well plates. Cells were detached using trypsin/EDTA and resuspended in DMEM supplemented with 0.4% FBS. A total of 5 × 105 cells was loaded in the upper well in a volume of 300 µl, and the lower well was filled with 750 µl of DMEM with 10% FBS. The plates were incubated at 37°C for 24h before removing the remaining cell suspension. The invasion insert was placed in a clean well containing 4% PFA for 1 h at room temperature, stained with crystal violet for 1 h, and washed three times in PBS. Cells on the upper side of the filters were removed with cotton-tipped swabs, and the number of migrated cells on the bottom side of the filter was counted in five randomly chosen fields at 200× magnification and averaged. All experiments were performed in triplicate, and each experiment was repeated at least three times.

##### *Anchorage-dependent colony formation assay*

Anchorage-dependent growth was monitored as described previously (26). Briefly, anchorage-dependent growth was monitored on solid (plastic) surface. Approximately 2,000 HeLa cells stably expressing WT, 3YF, or Y154H Gαi were plated in 6-well plates and incubated in 5% CO2 at 37°C for ~2 weeks in 2% FBS growth media. Colonies were then stained with 0.005% crystal violet for 1 hr. Each experiment was analyzed in triplicate.

### Supplementary Figures and Figure Legends

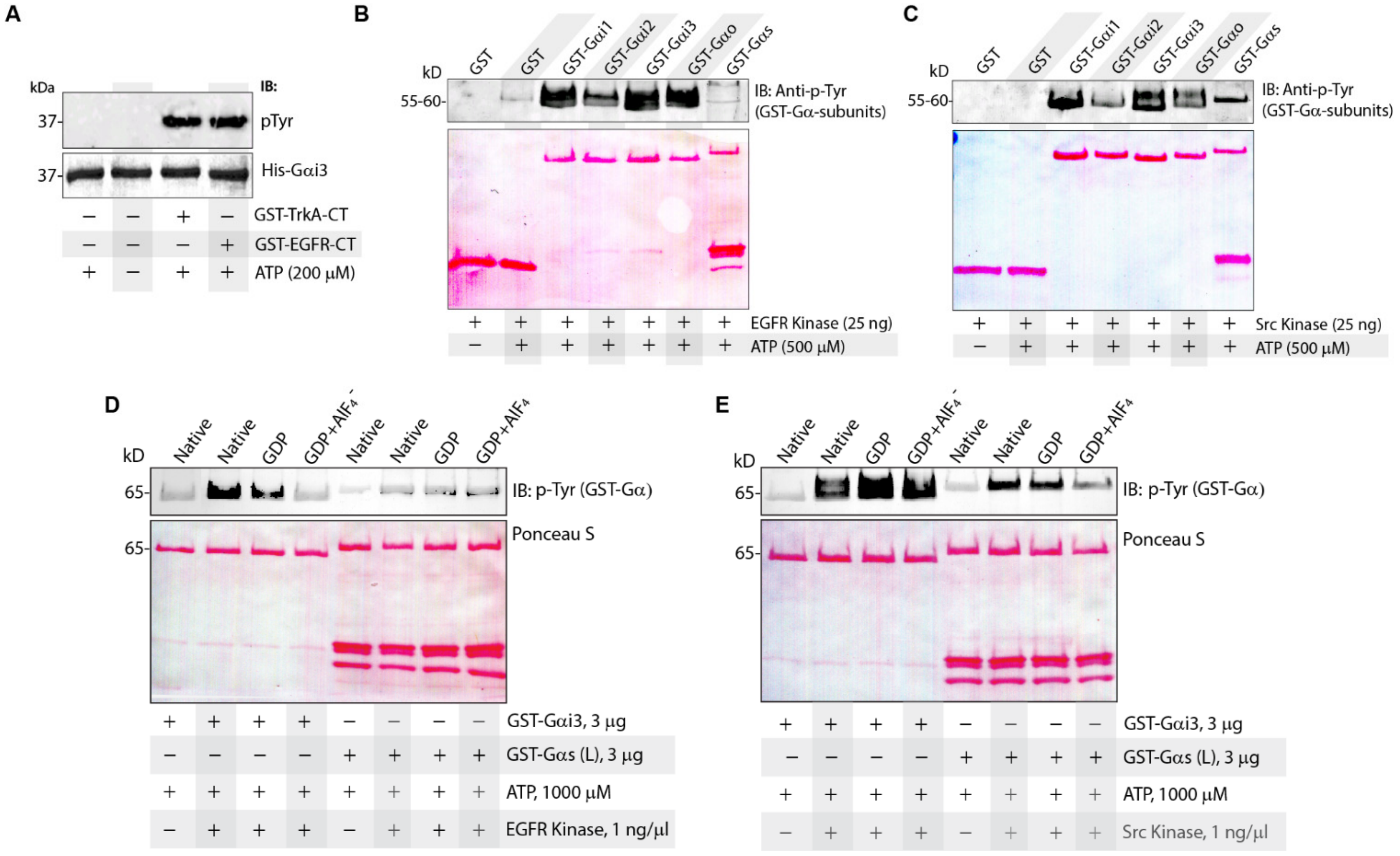


**Figure S1**: **Phosphorylation of Gαi/s subunits using recombinant active receptor and non-receptor (Src) tyrosine kinases. A**, In vitro kinase assays using recombinant active TrKA and EGFR as kinases and Gαi3 as substrate. Entire reactions were analyzed for the extent of phosphorylation by immunoblotting with anti-pTyr mAb. Similar amounts of substrate protein was confirmed by ponceau S staining. **B**, In vitro kinase assays using recombinant EGFR and different Gαi/o/s subunits as substrates were analyzed as in A. **C**, In vitro kinase assays using recombinant activated c-Src and different Gαi/o/s subunit as substrates were analyzed as in A. **D**, In vitro kinase assays using recombinant EGFR and Gαi3/Gαs subunits at native, inactive and active state as substrates were analyzed as in A. **E**, In vitro kinase assays using recombinant activated c-Src and Gαi3/Gαs subunits at native, inactive and active state as substrates were analyzed as in A.

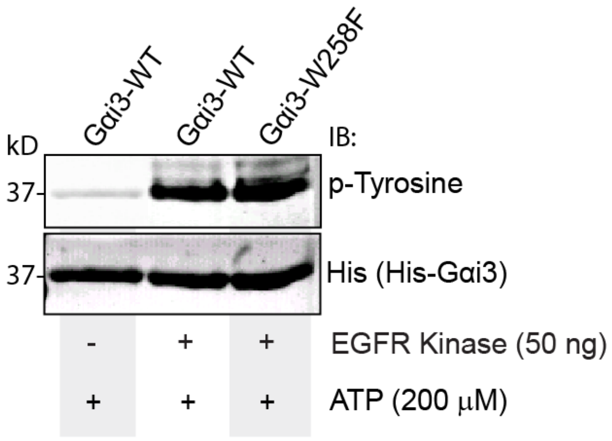


**Figure S2**: **Recombinant active EGFR can phosphorylate WT and W258F mutant Gαi proteins to similar extent *in vitro*.** *In vitro* kinase assay of WT or W258F mutant His-Gαi3 protein with EGFR. The entire reaction was analyzed by dual-color immunoblotting for tyrosine phosphorylated His-Gαi using anti-pTyr mAb and anti-His pAb.

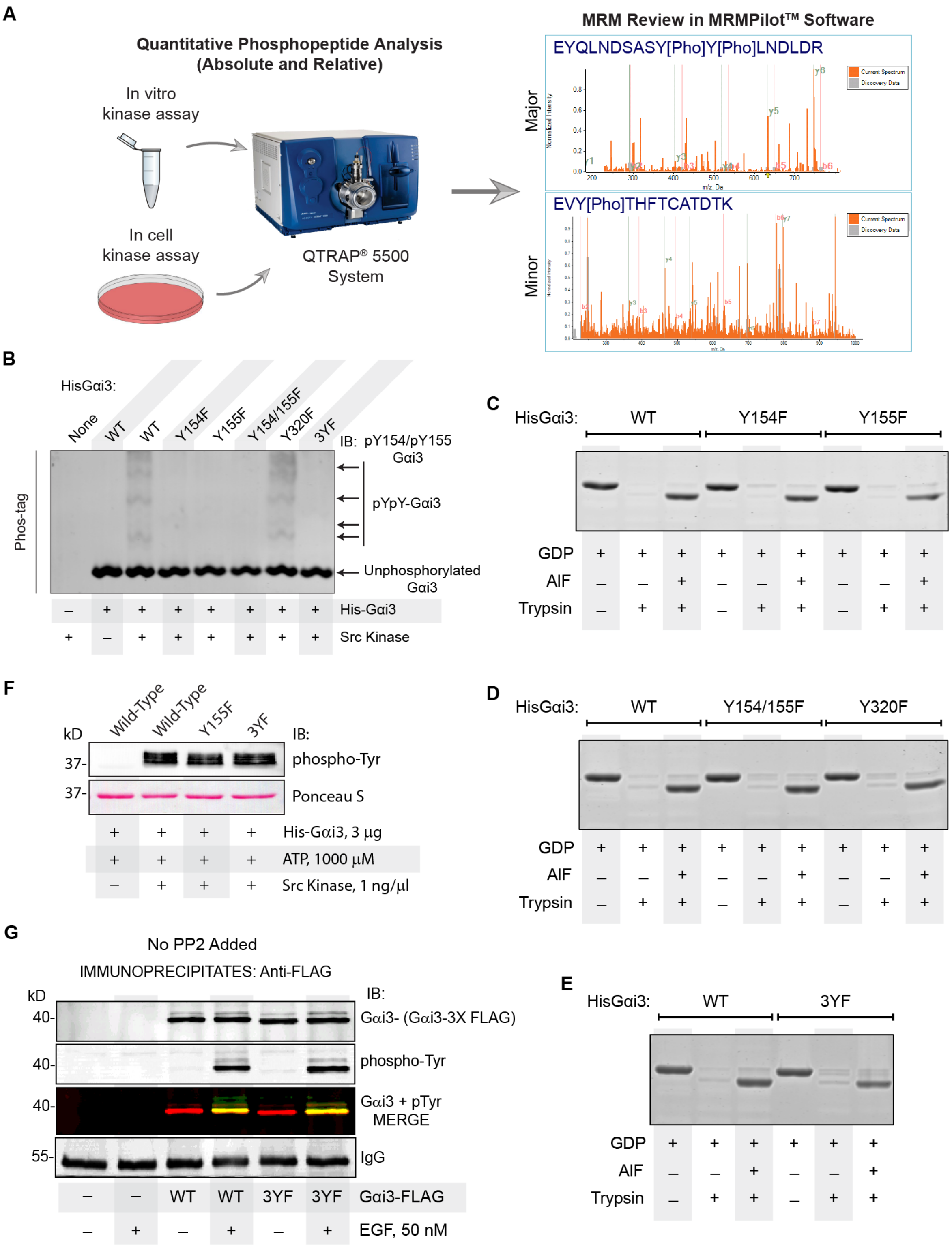


**Figure S3. RTKs directly phosphorylate Gαi on unique sites.**
**A**, Schematic displaying experiment approach for identification of RTK phosphorylation sites on Gαi3. **B**, In vitro kinase assay of Src with His-Gαi3-WT and various non-phosphorylaS YF mutants run on Phos-tag® gel and immunoblotted with custom pYpY-Gαi3 antibody. **C-E**, Coomassie stain of trypsin proteolysis assays performed on WT and tyrosine mutant Gαi3 proteins loaded with GDP or GDP-AlF$^{4}$ (latter mimics a GTP-bound-like state). As expected, the G protein is partially resistant to proteolysis in GTP-bound state (but not when GDP bound). **F**, In vitro kinase assays using recombinant active c-Src and WT or tyrosine mutant Gαi3 proteins as substrates were analyzed by immunoblotting with pan-pTyr antibody. **G**, Immunoprecipitation of transfected FLAG-tagged Gαi3-WT or Gαi3-3YF from EGF-stimulated Cos-7 cells in the absence of PP2 inhibitor were analyzed by immunoblotting with pan-pTyr antibody.

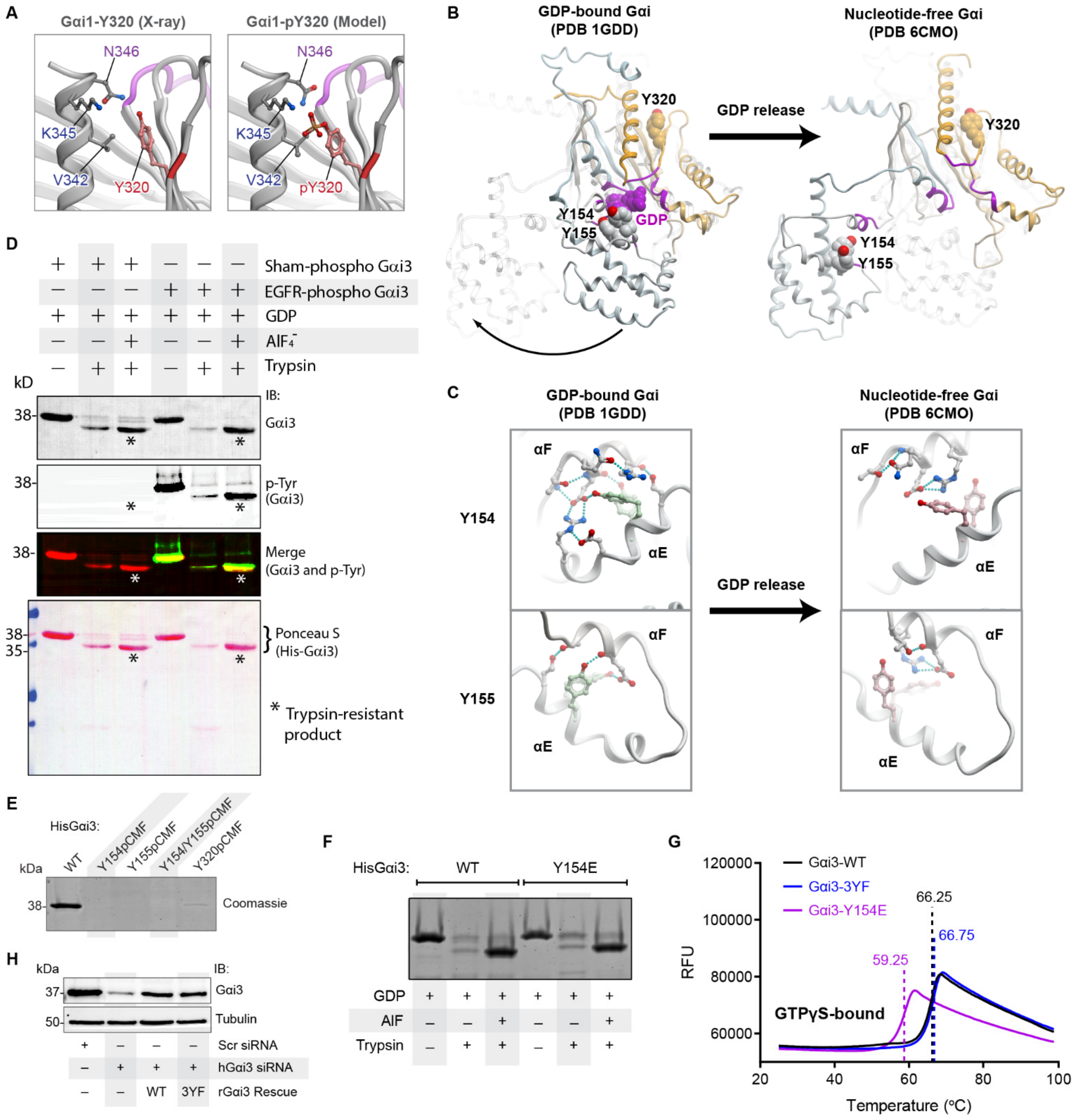


**Figure S4**: **Tyrosine phosphorylation of Gαi by RTKs may alter conformation, reduce stability. A**, Structures of Gαi1 highlighting Y320 and neighboring residues with and without being phosphorylated. **B**, Crystal structures of Gαi in the GDP-bound (*left*) and nucleotide-free (*right*) states, highlighting the positions of Y154, Y155 and Y320 in these structures. **C**, Crystal structures of Gαi showing close ups of the hydrogen-bond network made by Y154 and Y155 in the GDP-bound (*left*) and nucleotide-free (*right*) states. **D**, Trypsin proteolysis assays performed on in vitro ‘sham’ and EGFR phosphorylated WT Gαi3 proteins subsequently loaded with GDP or GDP+AlF$_4^-$ (the latter mimics GTP-bound conditions). **E**, Coomassie stain of attempts to purify pCMF-incorporated Gαi3 protein shows failure to express/purify significant amounts of proteins in which Y154 and/or Y155 are replaced with pY-mimic unnatural amino acid. **F**, Coomassie stain of trypsin proteolysis assays performed on Gαi3-WT and Gαi3-Y154E proteins loaded with GDP or GDP-AlF$^4$. **G**, WT and Tyr mutant Gαi proteins were subjected to increasing temperatures in differential scanning fluorimetry (thermal shift) assay. Findings are displayed as a line graph showing average RFU curves of GTPγS-bound (40µM GTPγS added). Measured melting temperature for each condition is indicated by the vertical dotted lines. **H**, Western blot validation of stable Gαi3-WT or Gαi3-3YF HeLa rescue cell lines.

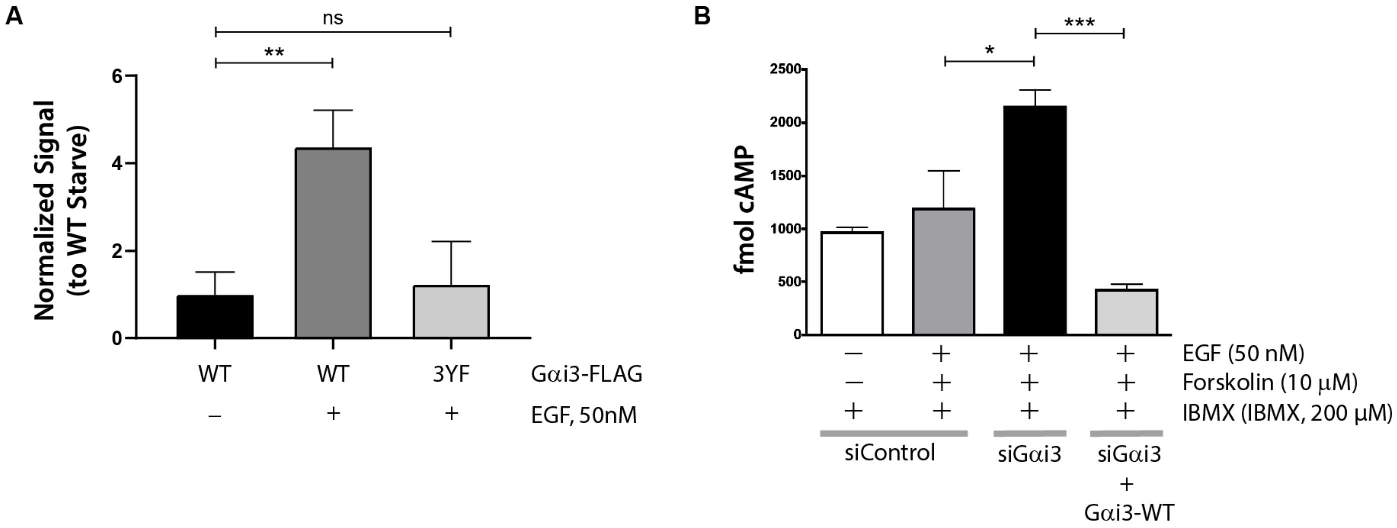


**Figure S5**: **Phosphorylation by EGFR activates Gαi in cells.**
**A**. Bar graphs showing the relative levels of activated FLAG-tagged Gαi3-WT or Gαi3-3YF in cells responding to EGF, as detected by immunoprecipitation using the Gαi:GTP (active conformation specific) antibody. Immunoblots are shown in **Fig. 4G**. **B**. Bar graphs showing the absolute levels of cAMP, as detected by radioimmunoassay (RIA) in TCA extracts of HeLa cells depleted (siGαi3) or not (siControl) of endogenous Gαi3 and then rescued by stably expressing rat Gαi3-WT. All cells were serum starved (0.2 % FBS, 16 h) and incubated with isobutylmethylxanthine (IBMX, 200 μM, 20 min) followed by EGF (50 nM, 10 min) and Forskolin (10 μM, 10 min), as indicated.

A

| GENE | UniProt ID | RefSeq ID | Ensembl Peptide | Chrom. | Chrom. Position | Ref Base | Alt Base | Protein Position | Ref Residue | Alt Residue | Cancer Type | # of Tumors | Source |
|---|---|---|---|---|---|---|---|---|---|---|---|---|---|
| **GNAI1** | P63096-1 | NP_002060.4 | ENSP00000343027 | 7 | 79,833,022 | A | C | 155 | Y | S | DOID:1324/ lung cancer | 2 | TCGA |
| **GNAI2** | P04899-1 | NP_002061.1 | ENSP00000460235 | 3 | 50,290,615 | T | C | 154 | Y | H | DOID:3571/ liver cancer | 2 | COSMIC |
| | | | | | | T | C | 154 | Y | H | DOID:0070003/ blastoma | 1 | ICGC |
| **GNAI3** | P08754-1 | NP_006487.1 | ENSP00000358867 | 1 | 110,121,982 | T | G | 154 | Y | D | DOID:2531/ hematologic cancer | 1 | ICGC |

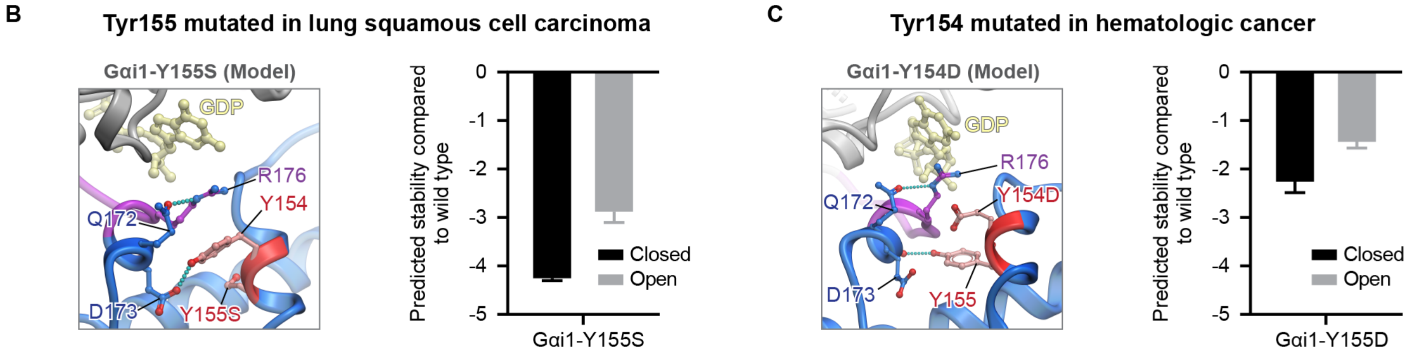


D

**Phospho-mimetic Gαi3 proteins attempted to be purified**

| Gαi3 Mutant Protein | Sufficient Yield for Biochemical Assays |
|---|---|
| Y154pCMF | No |
| Y155pCMF | No |
| Y154pCMF/Y155pCMF | No |
| Y154E | Yes |
| Y155E | No |
| Y154E/Y155E | No |
| Y154D | No |
| Y154H | Yes |
| Y155S | No |

**Figure S6**: **Somatic mutations targeting Y154/Y155 in tumors provide insights into how RTK-mediated tyrosine phosphorylation of Gαi affect cellular phenotypes. A**, Table summarizing the documented cancer mutations of Y154 and Y155 on all Gαi isoforms from three sources, the COSMIC, https://cancer.sanger.ac.uk/cosmic; cBioPortal, https://www.cbioportal.org/ and BioMuta using a High-performance Integrated Virtual Environment (HIVE) (27, 28). **B**, (*Left*) Structural model of Gαi1-Y155S cancer mutation highlighting hydrogen bonds. (*Right*) Bar graph displaying computationally predicted structural stability of Gαi1-Y155S in the open and closed states. **C**, (*Left*) Structural model of Gαi1-Y154D cancer mutation highlighting hydrogen bonds. (*Right*) Bar graph displaying computationally predicted structural stability of Gαi1-Y154D in the open and closed states. **D,** Table summarizing the purification attempts for phospho-mimetic and cancer mutant Gαi3 proteins and the yield from those attempts.

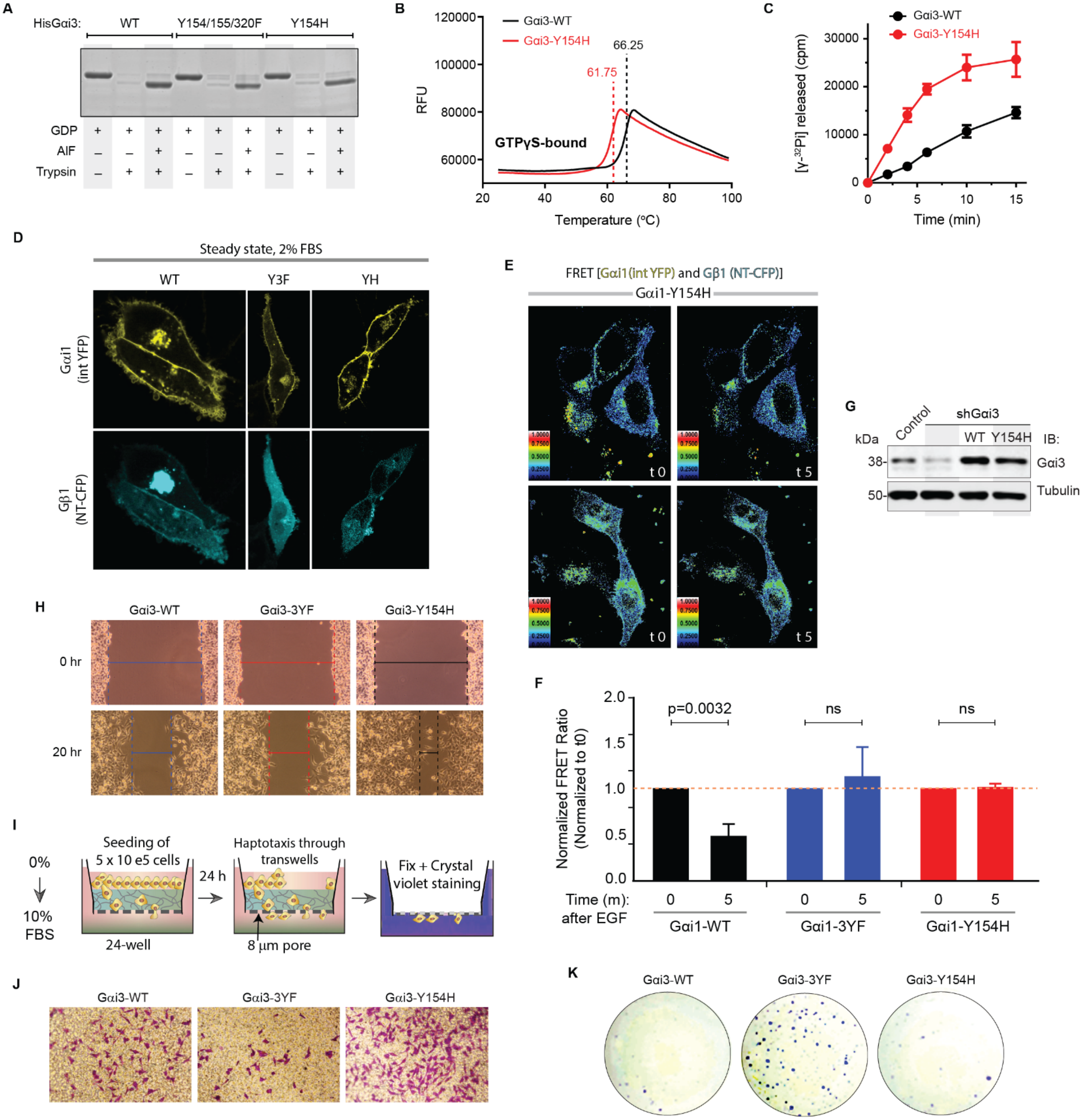


**Figure S7**: **Somatic mutations targeting Y154/Y155 in tumors provide insights into how RTK-mediated tyrosine phosphorylation of Gαi affect cellular phenotypes. A**, Coomassie stain of trypsin proteolysis assays performed on WT and tyrosine mutant Gαi3 proteins loaded with GDP or GDP-AlF$^4$. WT and 3YF data shown here are same as in Figure 3—figure supplement 1D. **B**, WT and Y154H mutant Gαi proteins were subjected to increasing temperatures in differential scanning fluorimetry (thermal shift) assay. Findings are displayed as a line graph showing average RFU curves of GTPγS-bound (40μM GTPγS added). Measured melting temperature for each condition is indicated by the vertical dotted lines. **C**, Dot plot displaying the GTPase activity of WT and Y154H mutant Gαi proteins determined by quantifying the amount of [$^{32}$P]Pi released from [γ-$^{32}$P]GTP at the indicated time points. Data (c.p.m.) are presented as mean ± SD of n=2. **D**, Representative CFP and YFP images of cells expressing Gαi1-Y154H activity reporter before and after EGF stimulation. Cells that expressed equimolar amounts of donor and acceptor probes were chosen selectively for FRET analyses. Images displayed are CFP and YFP channels from experiment shown in **Fig. 5F**. **E**, Representative FRET images of cells expressing Gαi1-Y154H activity reporter before and after EGF stimulation. FRET scale is shown in inset. **F**, Bar graphs displaying quantification of FRET results from (**E**). Error bars, ±S.E.M.; n = 5-7 cells/experiment, from 4 independent experiments. Gαi1-WT and Gαi1-3YF data are same as shown in **Fig. 4K**. Data for all mutants were collected side-by-side on the same day. **G**, Western blot validation of stable Gαi3-WT or Gαi3-Y154H HeLa rescue cell lines. **H**, Representative images of cell stably expressing WT or mutant Gαi proteins at different time points during 2D wound closure cell migration assays conducted under 2% FBS conditions. Quantifications are shown in **Fig. 5H**. **I**, Schematic showing workflow for 3D migration across serum gradient. **J**, Representative images showing cells stably expressing WT or mutant Gαi proteins that successfully migrated across the transwell within 24 h during 3D haptotaxis-transwell assays conducted using a 10% serum gradient. Quantifications are shown in **Fig. 5I**. **K**, Representative images of cell stably expressing WT or mutant Gαi proteins after 14 days of anchorage-dependent colony formation growth conducted under 2% FBS conditions. Quantifications are shown in **Fig. 5J**.

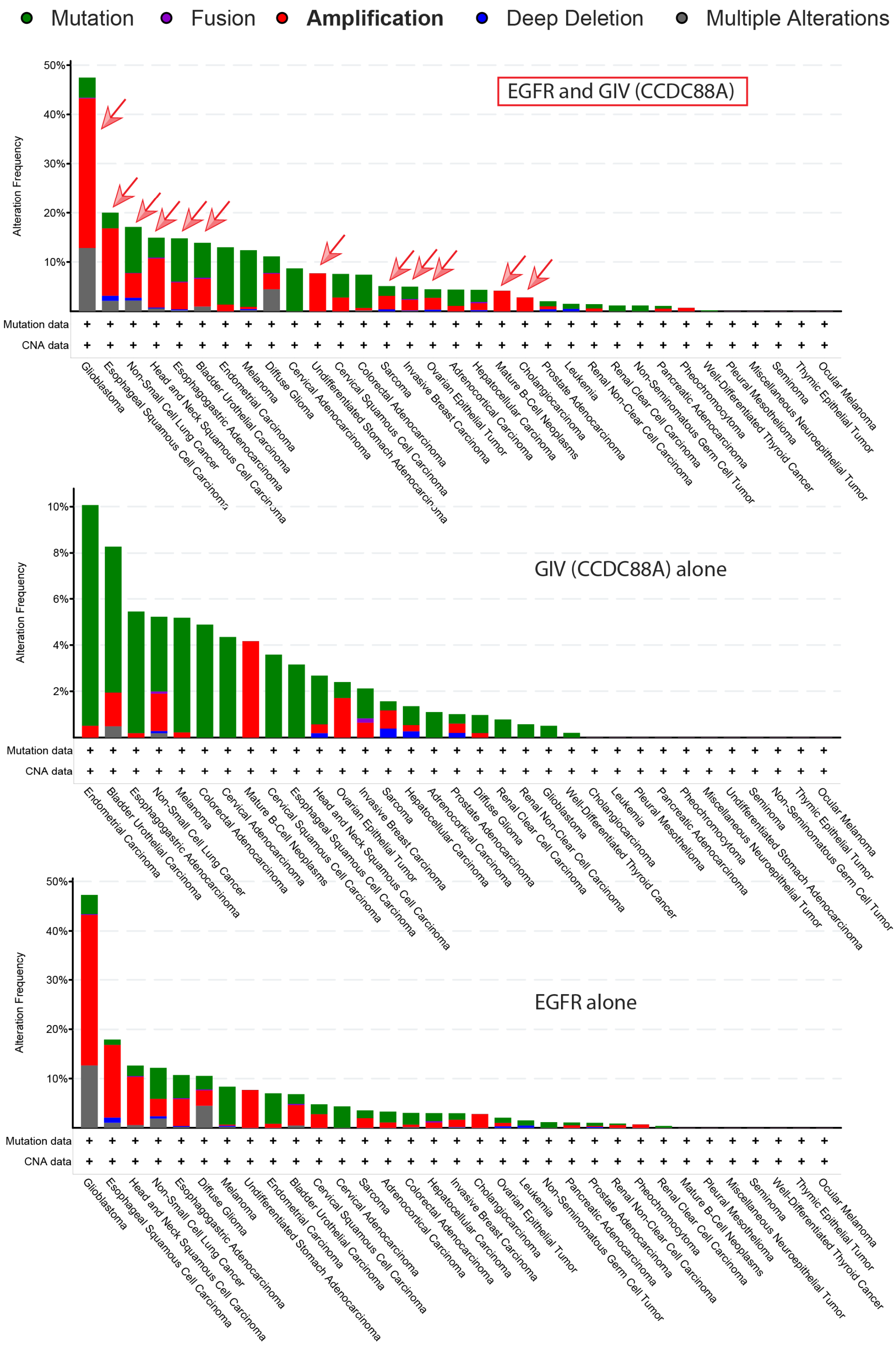


**Figure. S8**: **EGFR and GIV levels are frequently increased in different types of cancers.** A query of EGFR and CCDC88A (GIV) in pan-cancer TCGA datasets [10953 patients / 10967 samples in 32 studies] using cBioportal (cBioPortal.org) reveals the types of tumors where EGFR and/or GIV (top), or GIV alone (middle), or EGFR alone (bottom) is altered. Key on top shows the color code corresponding to the type of alteration. Co-occurrence of EGFR and GIV amplification is seen in several kinds of tumors (arrows, top panel).

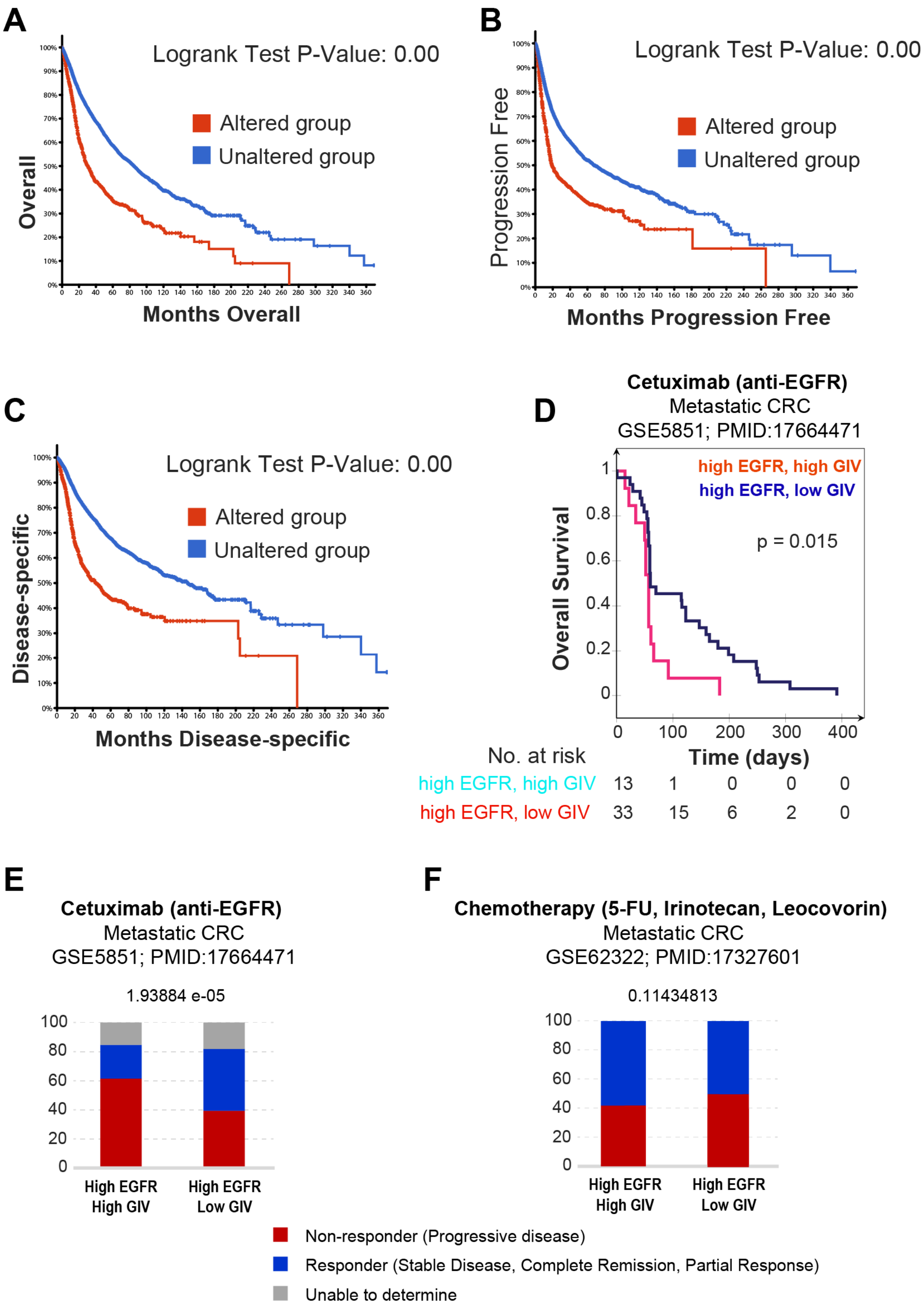


**Figure S9**: **Increased expression of EGFR and/or GIV carries poor prognosis and high expression of GIV in the setting of high EGFR is associated with resistance to anti-EGFR therapeutics. A-C,** Kaplan-Meier curves showing overall (**A**), progression-free (**B**) and disease-specific (**C**) survival rates stratified based on whether the levels of mRNA for EGFR and/or GIV were elevated in tumors of a pan-cancer TCGA dataset [10953 patients / 10967 samples in 32 studies] using cBioportal (cBioPortal.org)]. **D-E**, Kaplan-Meier curve show overall survival (**D**) and bar graphs display the treatment response status (**E**) in a cohort of patients with metastatic colorectal cancers receiving anti-EGFR targeted therapy (Cetuximab), stratified based on high *vs*. low GIV status. Statistical significance was estimated using Chi Sq. test (n = 80; GSE5851(29)) **F**, Bar graphs display the treatment response status in a cohort of patients with metastatic colorectal cancers receiving conventional combination chemotherapy, stratified into high *vs*. low GIV groups as in **E**. Statistical significance was estimated using Chi Sq. test (n = 180; GSE62322(30)).
